# Systematic Comparison of High-throughput Single-Cell and Single-Nucleus Transcriptomes during Cardiomyocyte Differentiation

**DOI:** 10.1101/585901

**Authors:** Alan Selewa, Ryan Dohn, Heather Eckart, Stephanie Lozano, Bingqing Xie, Eric Gauchat, Reem Elorbany, Katherine Rhodes, Jonathan Burnett, Yoav Gilad, Sebastian Pott, Anindita Basu

## Abstract

A comprehensive reference map of all cell types in the human body is necessary for improving our understanding of fundamental biological processes and in diagnosing and treating disease. High-throughput single-cell RNA sequencing techniques have emerged as powerful tools to identify and characterize cell types in complex and heterogeneous tissues. However, extracting intact cells from tissues and organs is often technically challenging or impossible, for example in heart or brain tissue. Single-nucleus RNA sequencing provides an alternative way to obtain transcriptome profiles of such tissues. To systematically assess the differences between high-throughput single-cell and single-nuclei RNA-seq approaches, we compared Drop-seq and DroNc-seq, two microfluidic-based 3’ RNA capture technologies that profile total cellular and nuclear RNA, respectively, during a time course experiment of human induced pluripotent stem cells (iPSCs) differentiating into cardiomyocytes. Clustering of time-series transcriptomes from Drop-seq and DroNc-seq revealed six distinct cell types, five of which were found in both techniques. Furthermore, single-cell trajectories reconstructed from both techniques reproduced expected differentiation dynamics. We then applied DroNc-seq to *postmortem* heart tissue to test its performance on heterogeneous human tissue samples. We compared the detected cell types from primary tissue with iPSC-derived cardiomyocytes profiled with DroNc-seq. Our data confirm that DroNc-seq yields similar results to Drop-seq on matched samples and can be successfully used to generate reference maps for the human cell atlas.

## Introduction

The identification and characterization of cell types from solid tissues and organs in the human body is the necessary basis for a comprehensive reference map of all human cells^1^. Such tissue atlases will provide a basis for understanding fundamental biological processes and to diagnose and treat disease. Single-cell RNA-sequencing (scRNA-seq) has emerged as a key tool to decompose complex tissues into cell types and states, and to investigate cellular heterogeneity^2–5^. Profiling cellular heterogeneity using thousands of cells and creating tissue level cellular maps require efficient and scalable scRNA-seq protocols. The development of microfluidic droplet-based approaches, such as Drop-seq, has enabled transcriptional profiling of thousands of cells in parallel^5,6^. Drop-seq has been used to characterize the cellular composition of a wide variety of tissues and organisms, including the mouse retina^5^, malaria parasites^7^, and drosophila embryos^8^. However, Drop-seq requires suspensions of intact single cells for library preparation which cannot be obtained for many tissues and cell types because of extra-cellular matrix that may be hard to digest, fragile cell membranes, unusual cell morphology, or large cell-size. This challenge may be addressed by adapting Drop-seq to single nuclei RNA-seq (DroNc-seq^9^). DroNc-seq obtains gene expression profiles from isolated nuclei which are more amenable for direct dissociation from tissues while maintaining membrane integrity. Both approaches can be used to characterize cellular composition of complex tissues. Comparisons of low-throughput, high-coverage single cell and single nucleus approaches suggest that both methods capture the cellular composition of heterogeneous samples to a similar degree^10,11^. However, direct comparisons of Drop-seq and DroNc-seq on matched samples have been limited to cell lines^9^ and, more recently, samples from mouse kidneys^12^. To establish a firm understanding of the differences and similarities of Drop-seq and DroNc-seq, it is necessary to compare these technologies across a spectrum of different biological conditions. A crucial aspect of single cell RNA-seq approaches is to capture cellular heterogeneity associated with expression changes during dynamic processes, for example during differentiation. We performed a systematic comparison of Drop-seq and DroNc-seq using time-course data from human iPSCs differentiating into cardiomyocytes (CMs). This allowed us to compare Drop-seq and DroNc-seq with respect to read depth, transcriptome composition, cell types detected, and cellular differentiation trajectories. These assessments are important for integrative analyses and interpretation of data produced using high-throughput single-cell and single-nucleus RNA-seq in general, and with Drop-seq and DroNc-seq in particular. In addition, we confirmed that inclusion of reads from intronic regions increases the sensitivity of DroNc-seq and improves resolution in identifying cell types. Next, we applied DroNc-seq to frozen *postmortem* human heart tissue to sample constituent cell types and compare them to CMs grown *in vitro* from human iPSC. This work was conceived as part of benchmarking experiments to establish the applicability of recent high-throughput single-nucleus RNA-seq for the Human Cell Atlas (HCA)^1^. By identifying differences and similarities between Drop-seq and DroNc-seq, this study will aid efforts such as the HCA that require the integration of single-cell and single-nucleus RNA-seq data from various tissues and laboratories into a common platform.

## Results

To quantitatively assess the similarities and differences in transcription profiles from single-cell and single-nucleus RNA-seq, we performed Drop-seq and DroNc-seq, respectively, on cells undergoing iPSC to CM differentiation, following an established protocol^13^. To compare Drop-seq and DroNc-seq across samples with different cellular characteristics and degrees of heterogeneity, we collected cells from multiple time-points throughout the differentiation process (days 0, 1, 3, 7, and 15) (Figure 1A). For each technique, we obtained samples from two cell lines per time-point, except for time-point day 15 which contains cells from a single cell line. DroNc-seq also contains a single cell line for day 1. To approximate how many cell barcodes were accidentally associated with 2 cells in our experiment (doublet rate), we mixed iPSCs from chimp into the Drop-seq run from cell line 1 on day 7. These data confirmed a low doublet rate (<5%) (Figure S1). The distributions of number of genes for each day of differentiation are shown in Figure 1B. Overall, Drop-seq shows a higher number of genes and transcripts detected compared with DroNc-seq, reflecting the greater abundance of transcripts in the intact cell, compared with the nucleus alone. For our analyses, we selected cells and nuclei with at least 400 and 300 detected genes (at least 1 UMI), respectively, and removed chimp cells from the day 7 sample. After filtering, the mean number of genes detected per cell and per nucleus are 962 and 553, and the mean number of UMI per cell, nucleus are 1474 and 721 for Drop-seq and DroNc-seq, respectively. Based on the above cut-offs, we detected a total of 25,475 cells and 17,229 nuclei across all cell lines and time-points for Drop-seq and DroNc-seq, respectively. Both cell lines were present at each time-point in the filtered datasets (Figure 1C). Using raw RNA-seq reads, we found that top expressed genes in Drop-seq comprised of mitochondrial and ribosomal genes, while the top gene in DroNc-seq was the non-coding RNA, MALAT1 (Figure 1D).

**Figure 1:**
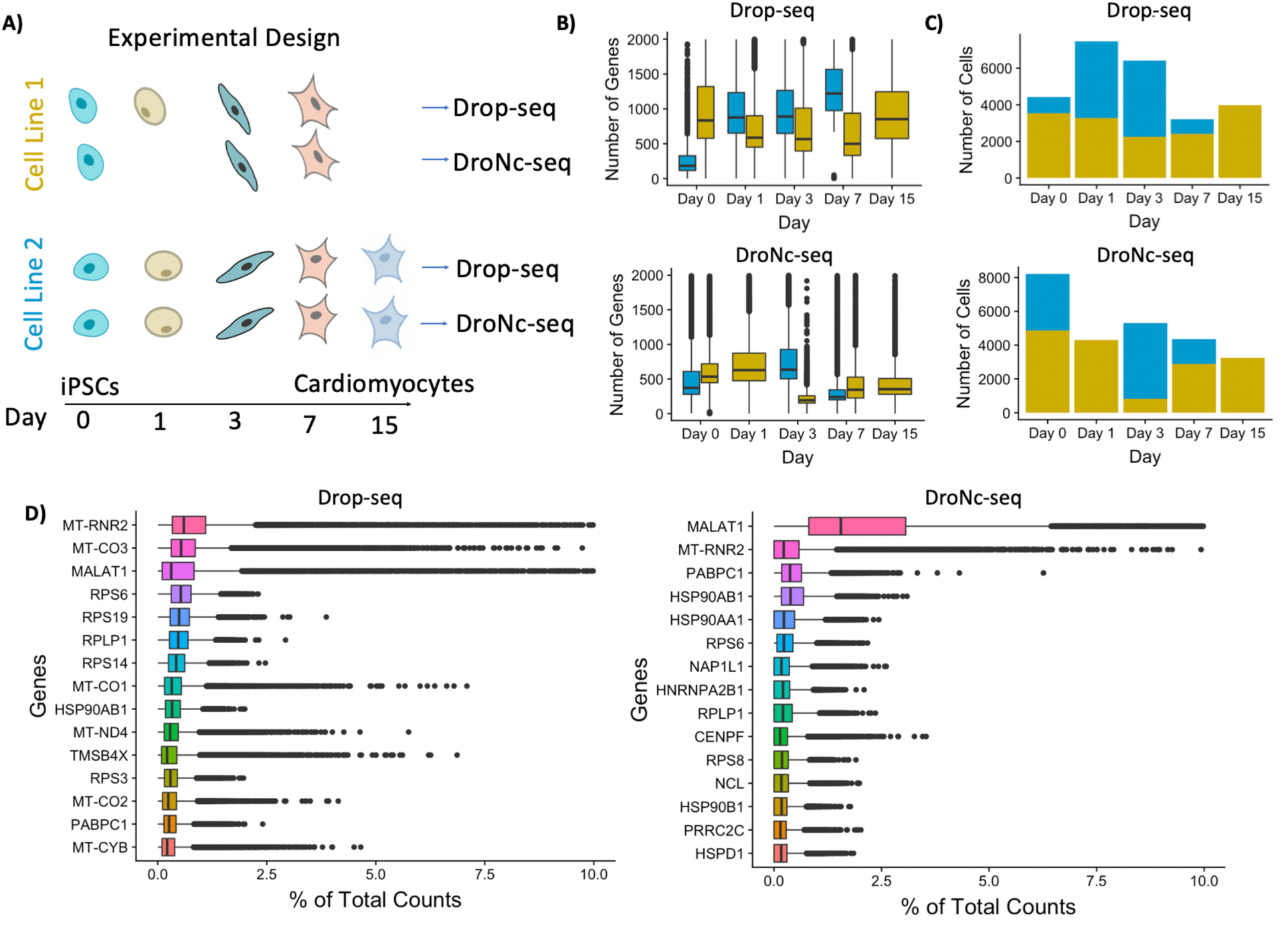
Experimental design and preliminary data analyses. A) Two cell lines of iPSCs differentiating into CMs over a 15-day time period underwent mRNA sequencing with Drop-seq and DroNc-seq. B) Boxplots showing the distribution of number of genes in each day and cell line for Drop-seq (top) and DroNc-seq (bottom). C) Number of cells present after applying quality control cut-offs. D) Percentage of counts for the top 15 genes in Drop-seq (left) and DroNc-seq (right).

In addition to the differences in the number of genes detected in Drop-seq and DroNc-seq, DroNc-seq captures a significantly higher fraction of intronic reads compared with Drop-seq (Figure 2A). Up to 50% of the reads from DroNc-seq mapped to intronic regions, while for Drop-seq, only 7% of reads were intronic. This discrepancy between the two techniques is expected and likely caused by the sampling of unprocessed transcripts that are enriched in the nucleus. Intronic reads will be detected if the transcript was not fully processed before capture by the polydT primer. In addition, internal priming^14^ on polyA stretches might lead to further sampling of introns. In order to understand the sources of intronic reads in our dataset, we scanned the genome for polyA stretches that are at least 5 bp long, and counted their frequency within and around each read with 20 bp flanking regions. We found that approximately 40% of the intronic reads and their 20-bp flanking regions contained at least one polyA stretches and that these polyA stretches were specifically enriched towards the 3’ end of reads (Figure S3). This suggests internal priming as a contributing mechanism for intronic read sampling. RNA-seq reads aligning to introns have been used to quantify gene expression levels previously^11,12^. Indeed, incorporating intronic reads to quantify gene expression level improves the gene detection rate in DroNc-seq by ∼2 times on average (Figure 2B). This increase in detection rate leads to recovery of gene expression for cells which would otherwise not be detected, as demonstrated by examples from mesoderm and cardiac genes (Figure 2C). These data suggest that inclusion of introns can be used to compensate for the smaller amount of nuclear RNA compared with whole cells. Accordingly, we incorporated intronic reads into our analysis pipeline to improve gene detection rates in DroNc-seq. After intron inclusion, we recovered 1.5 times more nuclei, bringing our total to 25,429 nuclei using a minimum of 300 genes detected per nucleus. In addition, the mean number of UMI per cell increased from 721 to 918, while the mean number of genes detected per cell increased from 553 to 672.

**Figure 2:**
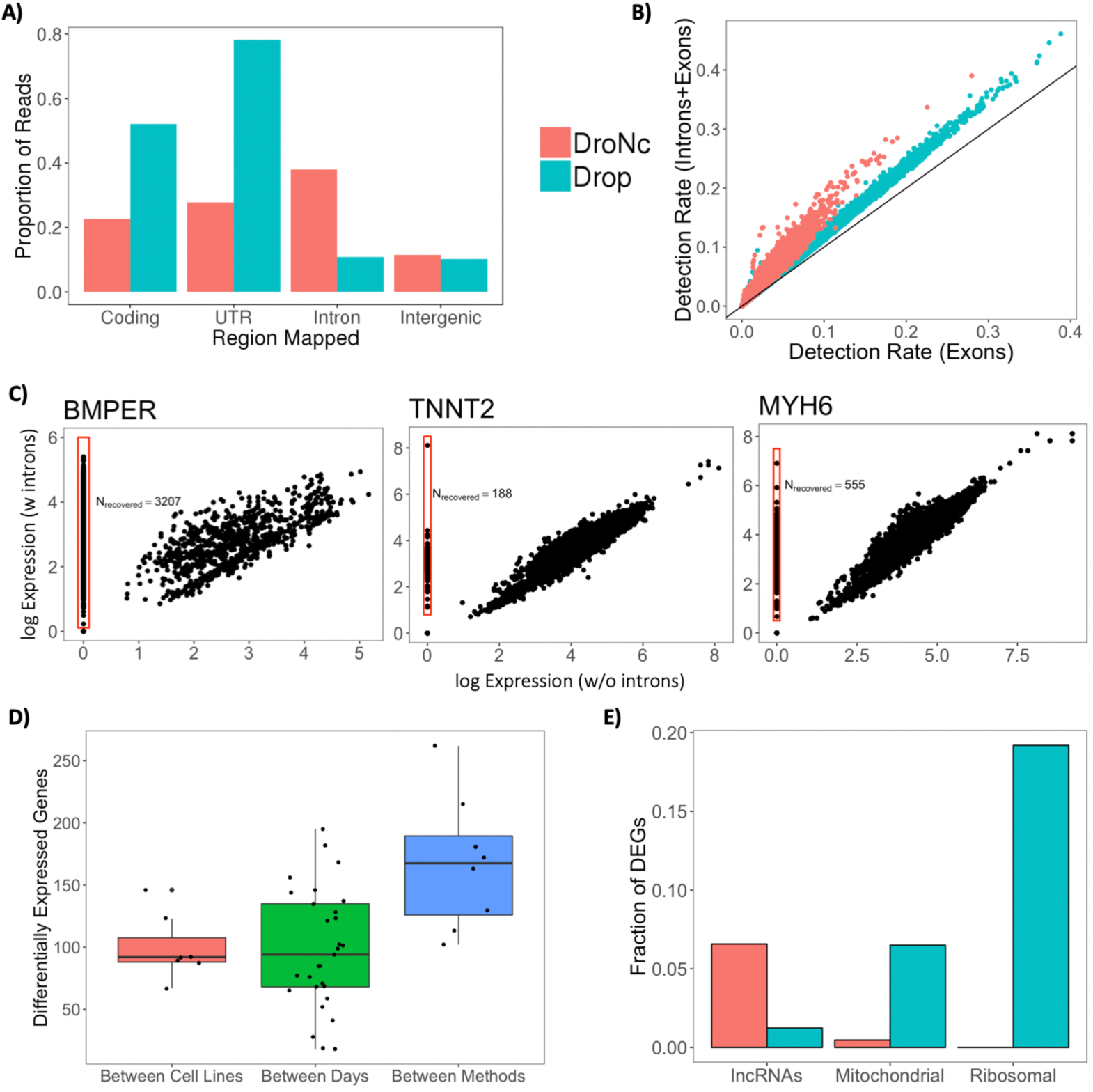
A) Distribution of reads across the genome in Drop-seq and DroNc-seq. B) Incorporating intronic reads in quantifying gene expression increases each cell’s gene detection rate by ∼2X on average for DroNc-seq, enabling detection of more genes per cell, compared with using exon reads only. C) Mesoderm and cardiac genes with expression detected when incorporating intronic reads. D) Differential expression analysis between methods, days, and cell lines. Genes with adjusted p-value < 0.05 and log-fold-change > 4 were kept. E) Proportion of differentially expressed genes (DEGs) between Drop-seq and DroNc-seq associated with different gene categories.

To identify systematic differences in gene-specific detection rates between Drop-seq and DroNc-seq, we obtained differentially expressed genes (DEGs) between the two techniques for matched time-points and cell lines. As a comparison, we also performed differential gene expression analyses between time-points and between cell lines within each technique. We detected substantially more genes with differential expression between the two techniques than we observed between different time-points or cell lines (Figure 2D). This phenomenon was most pronounced for highly significant genes and became less pronounced at more lenient thresholds of log fold-change (Figure S11). The differentially detected genes directly reflect the sampling differences in cellular components for the two techniques. GO analysis on DEGs between Drop-seq and DroNc-seq revealed functional annotations associated with the sampling of different cellular components of the two techniques (Figure S5). In particular, 5% of genes detected at higher levels in DroNc-seq were lncRNAs (compared to 1% in Drop-seq), while 20% and 6% of genes detected at higher levels in Drop-seq were mitochondrial and ribosomal transcripts, respectively (Figure 2E).

Next, we tested if the differences between Drop-seq and DroNc-seq in the number of detected UMI and enriched gene sets lead to inconsistent detection of cell types and variation in the inferred differentiation trajectory. To infer cell types found with Drop-seq and DroNc-seq data, we performed clustering of cells separately for each technique. We used the R package Seurat^15^ to perform normalization, dimensionality reduction, clustering, and visualization of individual cells, grouped by cell types (see Methods). Cell types were assigned to clusters based on comparison of genes that are significantly upregulated in the cluster to known marker genes. All genes were tested for differential expression using a negative binomial likelihood ratio test within the Seurat package and p-values were adjusted for multiple testing using Bonferroni correction. For each cluster, we ordered genes by their average log-fold-change (logFC) in descending order to identify marker genes, as genes associated with cell type have a large fold-change in expression. Note that p-values (raw and adjusted) for all marker genes are small (adjusted p < 10^-5^). We used the top marker genes for each cluster to identify cell type specific genes (Figures S6 and S7). We found that the clusters identified by Drop-seq and DroNc-seq captured the anticipated differentiation from iPSCs to CMs over the course of 7 days (Figure 3A and B, Supplemental Figure 4). The cluster formed by cells from early time-points day 0 and day 1 contained pluripotent stem cells (Figure 3A and B, ‘iPSC’, orange cluster), in agreement with the expression of characteristic markers such as DPPA4. Cells harvested on day 3 mostly formed a separate cluster (‘Cardiac progenitors’, green cluster) composed of cells expressing markers concordant with cardiac progenitors (e.g. expression of *EOMES* (logFC=1.08), a mesendoderm progenitor marker gene). For days 7 and 15 the clusters of cells profiled by Drop-seq and DroNc-seq showed slight differences and we detected four clusters in Drop-seq compared to three for DroNc-seq, indicating that Drop-seq might be more sensitive towards detection. Drop-seq and DroNc-seq identified three clusters of ostensibly similar cell types. Two of these clusters contained cells predominantly expressing markers of CMs, including *MYH6, TNNT2, MYL*, and *MYBPC3* (Figure 3A, cyan cluster, ‘Cardiomyocyte 1’ and blue cluster, ‘Cardiomyocyte 2’). We also detected a cell cluster that expressed cardiac markers alongside markers of other lineages (e.g. *FOXA2* and *TTR*, pink cluster, ‘Alternative lineage 1’). Drop-seq revealed an additional smaller cluster (purple, ‘Alternative lineage 2’, expression of *FLT1*) for which we did not find an equivalent cell population in DroNc-seq. These ‘Alternative lineage’ clusters might represent cells at intermediate stages, failures of differentiation, or differentiation towards alternative lineages. This heterogeneity and the detection of mesendodermal and endodermal cell populations, including endothelial cells, is in agreement with previous scRNA-seq data obtained during iPSC to cardiomyocyte differentiation^16^.

**Figure 3:**
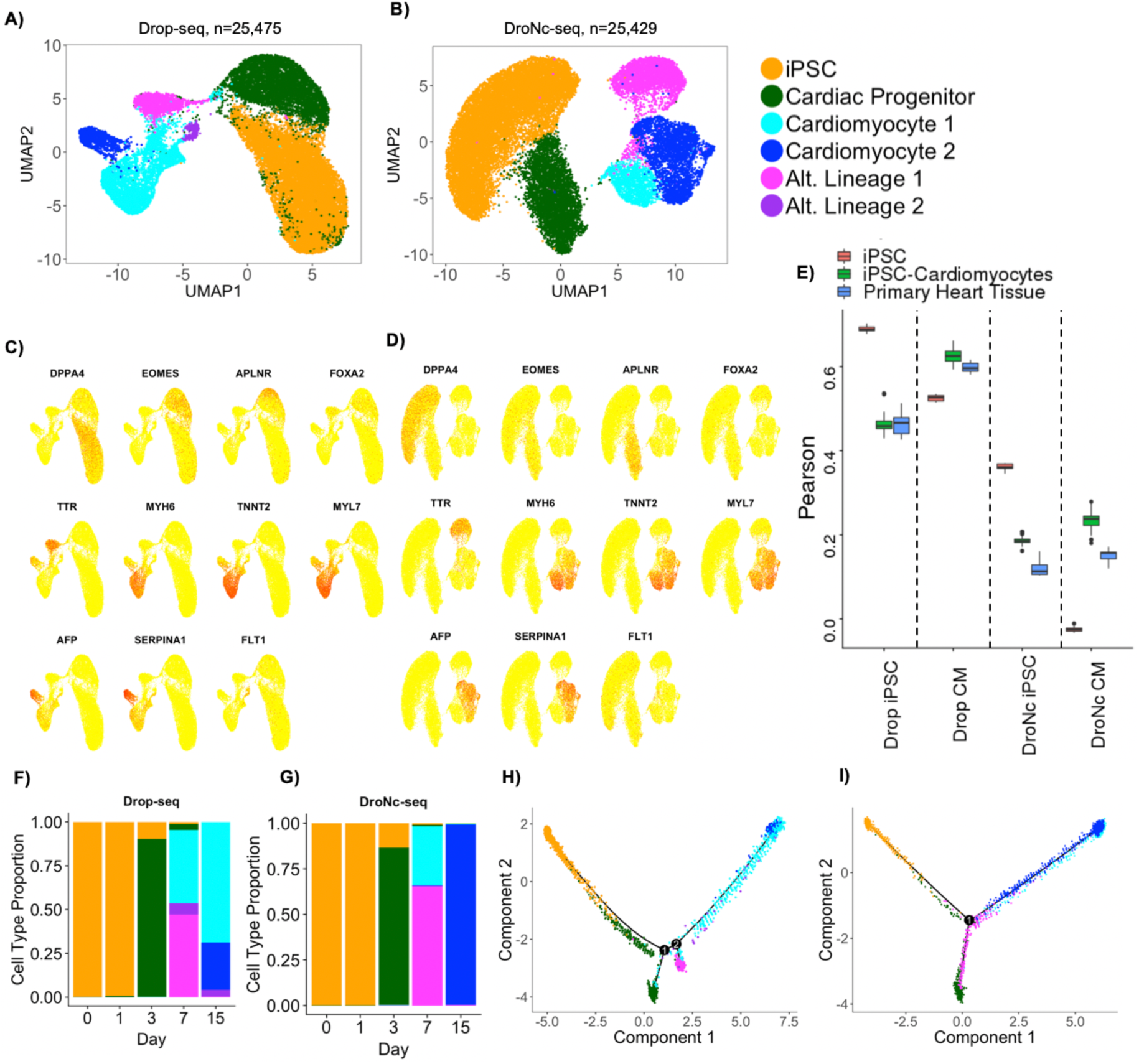
Cell type and single-cell trajectory analysis. A, B) Clustering results visualized with UMAP and colored by inferred cell type for Drop-seq and DroNc-seq. C, D) Expression of marker genes overlaid on UMAP plots from A and B for Drop-seq and DroNc-seq. E) Pearson correlation of DroNc-seq and Drop-seq pseudo-bulk against bulk RNA-seq from iPSCs (n=18), iPSC-Cardiomyocytes (n=51), and primary heart tissue (n=22)^17^. F, G) Distribution of cell types per time-point in Drop-seq and DroNc-seq, respectively. H, I) Inferred trajectories using Monocle with color representing inferred cell types. A total of 3500 cells were used for the trajectory corresponding to 700 per time-point.

**Figure 4:**
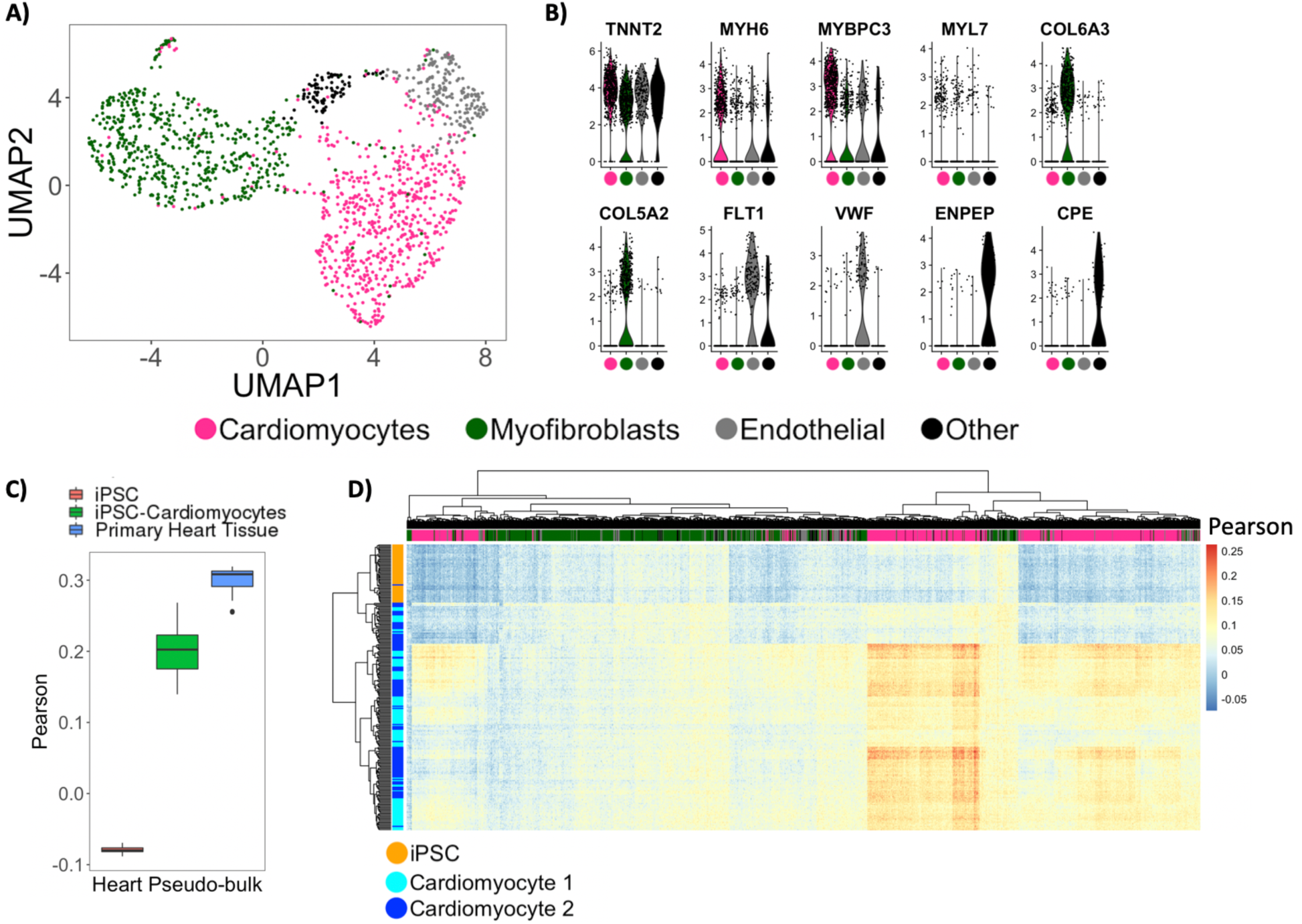
Application of DroNc-seq on human heart tissue. A) Cell type analysis visualized with UMAP. B) Distribution of marker genes identified with differential expression analysis. All genes listed have p-values < 10^-29^. C) Pearson correlation of primary heart pseudo-bulk against bulk RNA-seq from iPSCs (n=18), iPSC-Cardiomyocytes (n=51), and primary heart tissue (n=22)^17^. D) Bi-clustering on Pearson correlation values of primary heart nuclei with nuclei from iPSCs and iPSC-derived cardiomyocytes.

Table S1 shows the marker genes used to identify each cell type and its corresponding cellular prevalence. This comparison supported that both Drop-seq and DroNc-seq can identify the predominant cell types expected in a population. Importantly, the identified clusters showed expression of similar sets of genes in both techniques indicating that, despite differences in detection rate between the techniques and preferential detection of specific subsets of genes the identification of major cell types remained largely unaffected.

To test how concordant the cluster assignment of Drop-seq and DroNc-seq are with bulk RNA-seq of similar cell types, we aggregated clusters representing iPSCs and iPSCs-CMs into pseudo-bulk samples. We compared these pseudo-bulk data to bulk RNA-seq data obtained from a previous study^17^. A total of 91 bulk RNA-seq samples composed of human iPSCs (n=18), iPSCs differentiating into CMs (n=51), and adult primary heart tissue (n=22) were used for a correlation analysis against pseudo-bulk iPSCs and CMs (Figure 3E). Drop-seq generally outperforms DroNc-seq for all three sample types regardless of pseudo-bulk type by ∼ 50%, which is expected as bulk RNA-seq and Drop-seq both capture mRNA from whole cells. The iPSC pseudo-bulk samples of both methods are best correlated with iPSCs, followed by iPSC-Cardiomyocytes and primary heart tissue, as expected. For CM pseudo-bulk, both methods are best correlated with iPSC-cardiomyocytes, followed by primary heart tissue, and iPSCs.

The time-series data allowed us to compare differentiation dynamics of iPSCs captured by Drop-seq and DroNc-seq. We observed that several cell types were present in more than one time-point (Figures 3 F, G). In particular, iPSCs were observed in days 0 and 1, while CMs are observed in days 7 and 15 in both Drop-seq and DroNc-seq data. Detection of the same or similar cell types across time-points should therefore enable us to reconstruct continuous single-cell differentiation trajectories^14,18,19^ in an unsupervised manner to characterize the temporal relationship between different cell populations. Accordingly, we reconstructed differentiation trajectories of the cells from DroNc-seq and Drop-seq data using Monocle^19^. In order to reduce computational time, we selected the top 700 cells based on the number of genes detected at each time-point, for a total of 3,500 cells and used them to reconstruct the single-cell trajectory during iPSC to CM differentiation.

Inferred trajectories from DroNc-seq and Drop-seq data show one and two branching points, respectively. Coloring cells by cell type (Figures 3 H, I) and pseudo-time (Figure S9) confirms the temporal order of cell types in Figures 3 F, G. Monocle places iPSCs at the beginning of the trajectory, which has pseudo-time zero, followed by cardiac progenitors. Following cardiac progenitors along the trajectory, we find one branching point in DroNc-seq which broadly partitions CMs and the clusters associated with less well-defined cell types that might represent alternative lineage decisions or incomplete differentiation (Figure 3). In Drop-seq, these immature cells are on different branches and are both separated from the third branch containing CMs. These differences might reflect the higher gene expression fold differences observed for the genes we used to build the trajectories in Drop-seq compared to DroNc-seq. This might be a consequence of the lower read depth observed for DroNc-seq. Both methods suggested the differentiation of iPSCs into an intermediate cell type (cardiac progenitors), and finally a population of clearly identifiable cardiomyocytes, based on the expression of TNNT2 and MYH6, and a divergent trajectory towards alternative cell populations.

The comparison of Drop-seq and DroNc-seq data was motivated by the fact that Drop-seq cannot be applied to generate single-cell RNA-seq data from adult primary heart tissue, but DroNc-seq potentially can. Having established that DroNc-seq provides data ostensibly similar to Drop-seq in our *in vitro* setup, we applied DroNc-seq to frozen human heart tissue to identify possible cardiac cell sub-types and non-cardiac cells within the tissue.

We detected a total of 4,796 nuclei based on the presence of distinct cell barcodes using DroNc-seq on tissue from an adult human male heart. We used both introns and exons to quantify number of reads per nucleus, with mean number of genes and UMIs as 361 and 823, respectively. To focus our analyses on good quality nuclei, our analyses used the top 30% (1,491) of cells based on the number of genes detected. We performed cell type analysis on the heart cells using the same procedure as described for the *in vitro* samples. As expected, the majority of cells (∼82%) were CMs and myofibroblasts (Fig. 4A). Figure 4B shows the distribution of marker genes for each cell type obtained using negative binomial likelihood ratio test. A cluster was identified as CMs (Figure 4A, pink cluster) based on marker genes *TNNT2* (logFC=0.71), *MYH6* (logFC=0.87), and *MYBPC3* (logFC=1.38). A second cluster was identified as likely myofibroblasts (Figure 4A, dark-green cluster) expressing the collagen genes *COL5A2* (logFC=1.95) and *COL6A3* (logFC=1.92) and periostin (POSTN). Finally, a third cluster was identified as endothelial cells (Figure 4A, grey cluster) based on vascular endothelial growth factor receptor *FLT1* (logFC=2.4) and blood clotting protein *VWF* (logFC=1.77). A fourth clusr expressing *CPE* (logFC=2.4) and *ENPEP* (logFC=2.5) was identified likely representing myofibroblasts (Figure 4A, black cluster). Additional marker genes are listed in Figure S10, which shows the top 10 upregulated genes in terms of logFC in each cluster.

To better understand the cell type composition of the primary heart tissue, we first aggregated data from all nuclei into a pseudo-bulk heart sample and compared these to bulk data from iPSCs, iPSC-CMs, and primary heart tissue as before. We found that the pseudo-bulk heart sample most closely correlated with bulk RNA-seq data obtained from primary hearts, followed by iPSC-CMs. No correlation was observed with bulk iPSCs (Figure 4C). Second, to compare the heart nuclei data with the *in vitro* model we compared single nuclei of the heart to the DroNc-seq on iPSC-CMs using correlation analysis. Figure 4D shows a bi-clustered heatmap of the Pearson correlation coefficients with columns representing primary heart nuclei, and rows representing iPSC-CMs nuclei. Interestingly, hierarchical clustering of each (heart) nuclei’s Pearson values (columns) confirms the clustering pattern found in Figure 4A, which demonstrates the presence of non-cardiomyocytes within the single-nuclei primary heart sample. In particular, the cluster identified as CMs (Figure 4A, pink cluster) has stronger correlation values with the iPSC-CMs than the myofibroblasts and endothelial cells (Figure 4A, dark-green and grey clusters). Clustering the rows also revealed the relative correlation strengths of the two iPSC-CMs clusters with the primary heart nuclei. In particular, the ‘Cardiomyocyte 2’ cluster generally has stronger correlation with the primary heart nuclei than the ‘Cardiomyocyte 1’ cluster. This is could potentially reflect the observation that ‘Cardiomyocyte 2’ was associated with cells collected on day of our differentiation protocol and therefore to closer towards the mature state of CMs. We used iPSCs as an out-group for which we expect no correlation with primary heart nuclei, which is observed to be the case (Figure 4D).

## Discussion

Building a cell atlas of the human body requires the expression profiling of all human tissues from a range of different samples, including tissues that are hard to dissociate, composed of fragile cells, and frozen specimens, all of which are incompatible with single-cell RNA sequencing. As an alternative, DroNc-seq, a high-throughput single-nucleus RNA sequencing protocol, has the potential to reveal tissue heterogeneity, at scale, based on *nuclear* RNA, and is being increasingly used to profile primary tissue at high throughput. However, it is unclear how DroNc-seq compares with earlier single-cell RNA-seq protocols like Drop-seq across a range of different cell types and tissues. Previous studies have performed cell type comparisons using nuclear vs. whole-cell RNA using full-length mRNA sequencing assays at low throughput^10,11^. Drop-seq and DroNc-seq have been compared using adult mouse kidneys cells^12^. We performed a direct comparison of high-throughput, single-cell (Drop-seq) and single-nucleus (DroNc-seq) RNA-seq using iPSCs differentiating into CMs. Together with single-nucleus profiling of primary CMs from adult human heart tissue, this study enabled us to compare cell type detection, transcriptome profiling and infer cellular differentiation with two complementary high-throughput techniques, using an *in vitro* model of CM differentiation, and compare them directly to human primary CMs obtained from a frozen heart sample (see Methods) using DroNc-seq.

As expected, the number of UMIs per nucleus in DroNc-seq are lower than those for cells in Drop-seq. Consequently, the gene detection rate in DroNc-seq was significantly lower than for Drop-seq (Figure 1C). However, given the high number of reads in DroNc-seq that mapped to intronic regions we reasoned that inclusion of such reads might increase the gene detection rate. Indeed, intron inclusion significantly increased the sensitivity of DroNc-seq and improved cluster separation and cell type identification, in agreement with previous studies^10–12^. We also found that the inclusion of introns increased gene detection rate in single nuclei samples. Of note, a significant proportion of the intronic reads seems to originate not from transcripts primed at the 3’ end but from direct priming to polyA stretches in introns^14^ (Figure 2). While such reads still scale with the expression level of a transcript, the assumption that transcript levels are uniquely quantified by a single UMI may be violated in these cases.

Given the difference in input material, i.e., cellular vs. nuclear RNA, it is not surprising that we found a significant proportion of genes that are differentially expressed between Drop-seq and DroNc-seq samples. Some of the most highly enriched sets of genes reflected the technical differences between the two technologies. Genes specifically enriched in Drop-seq are ribosomal and mitochondrial. DroNc-seq presumably loses these transcripts that are predominantly localized in the cytoplasm. Conversely, as a class, lncRNAs are enriched in DroNc-seq which agrees with the nuclear localization of many of them.

Expression profiles in Drop-seq and DroNc-seq confirmed the differentiation of iPSCs into CMs and revealed major cell types found within the *in vitro* differentiation model of iPSC-CMs. These data also confirmed heterogeneity observed during differentiation. Drop-seq and DroNc-seq detected a population of cardiac progenitors with cellular prevalence 23.3% and 18.2%, respectively. They also both detected two clusters representing CMs: cardiomyocyte 1 (16.1% and 5.6% prevalence) and cardiomyocyte 2 (4.2% and 12.7% prevalence). Both methods also revealed a population of cells, ‘Alternative lineage 1’, that might represent alternative fate or that failed to reprogram fully, which accounted for 5.9% and 11.3% of all cells in Drop-seq and DroNc-seq, respectively. The presence of non-CMs during late-stage is expected for the *in vitro* differentiation model and has been observed previously^16^. Accordingly, the proportion of cells differentiating into CMs expressing TNNT2, assessed by FACS, varies widely between 20-80%^13^. Based on our cell type assignment in Drop-seq data, we obtained 28% and 29% cardiomyocytes on day 7 for the two cell lines and 70% CMs on day 15 for cell line 2, which fall within the expected range.

Drop-seq revealed an additional smaller cluster (purple, ‘Alternative lineage 2’, expression of *FLT1* and comprising 1.4% of the total population) for which we did not find an equivalent cell population in DroNc-seq. The reasons behind the failure of DroNc-seq to identify the small fraction of cells identified as ‘Alternative lineage 2’ in Drop-seq may be due to the lower capture rate of DroNc-seq (mean number of detected genes was 672) compared to Drop-Seq (mean number of detected genes was 962) (Figure S8) which might result failure of the clustering approach to resolve this sub-population in DroNc-seq, or due to the preferential loss of the particular cell type arising from DroNc-seq’s nuclei dissociation protocol. The mean number of genes detected in this subpopulation in Drop-seq was 1032, representing the cluster with the highest gene detection rate. It is possible that this facilitated the detection of this cluster in Drop-seq while the lower detection rate in DroNc-seq combined with the small number of cells corresponding to this cluster in the sample lead to the loss of this population during clustering. However, we cannot rule out specific loss or selection biases for of the cell type introduced during DroNc-seq sample preparation.

We chose the iPSC-to-CM differentiation because in addition to cell type detection, the highly heterogenous but temporally coordinated process allowed us to compare cellular lineages inferred based on Drop-seq and DroNc-seq data, respectively. Indeed, we were able to infer similar trajectories for both Drop-seq and DroNc-seq (Figure 3H and I). Both trajectories show continuous differentiation of iPSCs into cardiac progenitors along a single path, which then branches into CM and non-cardiac cells (progenitor cells and alternative lineages). This suggests that a substantial proportion of cells identified as CM progenitors in our cluster analysis are diverging from the differentiation trajectory relatively early on and ultimately are not becoming mature cardiomyocytes^16^. In the case of Drop-seq ‘Alternative lineage 1’ and ‘Cardiac progenitor’ cells are branching off on two separate points, while for DroNc-seq both populations are on one branch. The additional branching point might reflect the higher resolution achieved by Drop-seq.

Compared with bulk samples, Drop-seq pseudo-bulk is closer to tissue-level expression than DroNc-seq. This is expected as the tissue data represents RNA-seq data generated using whole cells, rather than nuclei. However, this difference does not mask cell type specific differences in the degree of correlation with bulk samples from iPSCs, iPSC-CM, and heart. Both Drop-seq and DroNc-seq CM pseudo-bulk correlate the best with bulk iPSC-CMs samples followed by primary heart tissue and iPSCs. While the iPSCs correlate best with the bulk iPSCs for both methods. The comparison with bulk samples provides further evidence for the cell type labels that were assigned based on marker genes.

Having demonstrated that Drop-seq and DroNc-seq performed similarly in detecting heart-like cell types, we applied DroNc-seq to primary heart tissue from adult human male. As expected, cell type analysis of the tissue revealed mostly CMs (43%) and (myo)fibroblasts (39%), as well as a smaller population of endothelial cells (12%). Interestingly, TNNT2 was detected in all the cell types but was significantly upregulated in the CM cluster. TNNT2 being a marker gene for CMs suggested the possibility that all nuclei are of the same broad category of cell type. Correlating transcription profiles from primary heart nuclei with the iPSC-derived CM nuclei further supports the inferred cell types from the primary heart tissue. The transcriptome profiles of primary heart nuclei that were assigned to the ‘Cardiomyocyte’ cluster are more strongly correlated with the profile of iPSC-CMs compared with primary heart nuclei in other clusters.

Sequencing of additional cells and increased read depth will help to increase the resolution and potentially lead to detection of additional cell types. However, it is important to keep in mind that tissue samples are not uniform mixtures of cell types. Thus, the creation of comprehensive cell maps likely requires sampling of a given tissue in multiple different locations, as seen from the relatively low cell type complexity in DroNc-seq data on the human heart tissue when sampled from only one anatomical region.

This comparison of Drop-seq and DroNc-seq demonstrates the capability of DroNc-seq in dissecting the multicellular environment within complex tissue such as the heart, which would otherwise not be possible with Drop-seq. We expect that DroNc-seq will be used to perform high-throughput transcriptomic profiling of tissues for which it is difficult to obtain suspensions of intact single cells and aid in initiatives such as the Human Cell Atlas and the Human Tumor Atlas.

## Methods

### Cell Culture and Differentiation

We used iPSCs from two individuals from a previously established panel of LCL-derived iPSCs^20^. iPSCs were seeded on 100 mm dishes 3-5 days prior to differentiation. At 70-100% confluency, growth media was replaced with heart media: RPMI (Thermo Fisher Scientific, 14-040-CM) supplemented with B-27 Supplement minus insulin (Thermo Fisher Scientific, A1895601), 2 mM GlutaMAX (Thermo Fisher Scientific, 35050-061), and 100 mg/mL Penicillin/Streptomycin (Corning, 30002Cl). A heart medium/Matrigel mix was made using this medium along with a 1:100 dilution of Matrigel (Corning, 35427) and 12 uM of the GSK-3 inhibitor CHIR99021 trihydrochloride (Tocris, 4953). This medium was changed to base heart media 24 hours later (Day 1). On Day 3, the previously described medium was replaced with heart medium containing 2 μM Wnt-C59 (Tocris, 5148). On days 5, 7, 10, 12 and 14 of the differentiation, media was refreshed with base heart media. Heart medium changes occurred daily. Beating CMs cells were observed around Day 7.

### Cell Processing

At each time-point, cells were harvested from 100 mm plates by treating with Accutase (BD Biosciences, #561527) to generate a single cell suspension; from Day 7 onward, a cell scraper was also employed to release adherent cells from plates. Cells were centrifuged at 300 xg for 5 minutes and supernatant was aspirated off. Cells were washed 3 times with 1X PBS, 0.01% BSA (NEB, #B9000S), henceforth called PBS-BSA). 10 μL of cells was combined with trypan blue for counting in an NI hemocytometer (InCyto, DHC-N01-2). Viability of cells at each time point was recorded (see Table 1). Cells were also labelled with a combination of 4′,6-diamidino-2-phenylindole or DAPI (Sigma, Cat #D9542) and Wheat Germ Agglutinin (WGA; Thermo Fisher Scientific, W11262) to assess nucleus and cell membrane integrity under fluorescence imaging, as shown in Figure 5A. 400,000 cells were taken and suspended in 2 mL PBS-BSA (200,000 cells/mL) for Drop-seq, and the remaining cells were used for nuclei isolation for DroNc-seq.

**Table 1:** Viability of harvested cells from each iPSC-CMs differentiation time-point

| Time Point | Date | Viability |
| --- | --- | --- |
| Time Course 1 Day 0 | 11/16/2017 | 70% |
| Time Course 1 Day 1 | 11/15/2017 | 50% |
| Time Course 1 Day 3 | 11/17/2017 | 80% |
| Time Course 1 Day 7 | 11/21/2017 | 60% |
| Time Course 2 Day 0 | 1/22/2018 | 60% |
| Time Course 2 Day 1 | 1/23/2018 | 80% |
| Time Course 2 Day 3 | 1/25/2018 | 80% |
| Time Course 2 Day 7 | 1/29/2018 | 90% |
| Time Course 2 Day 15 | 2/6/2018 | 55% |

**Figure 5:**
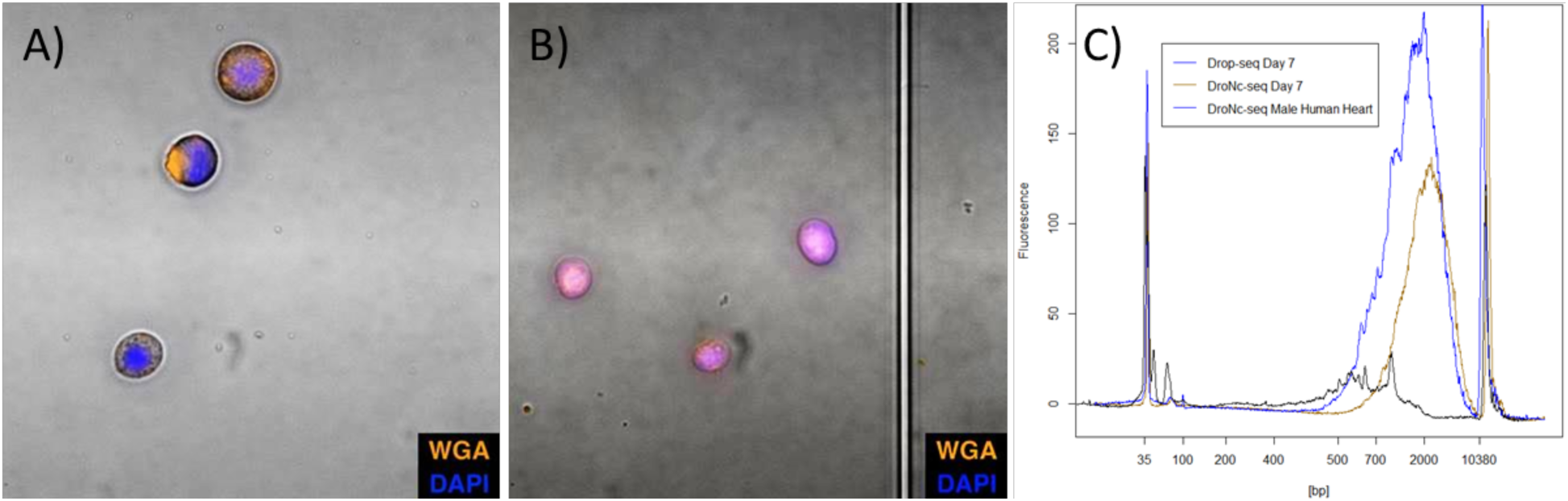
Experimental quality control metrics. Images of Day 1 of differentiation of human iPSC derived cardiomyocyte (iPSC-CM) cells-A) and nuclei-B) stained with DAPI and WGA; C) BioAnalyzer traces of WTA product from Drop-seq on iPSC-CM Day 7, DroNc-seq on iPSC-CM Day 7, and DroNc-seq on archived adult male heart tissue.

Nuclei were isolated using the Nuclei EZ Prep isolation kit (Sigma, Cat #NUC-101). Briefly, cells were resuspended in 4 mL EZ Prep Lysis Buffer and incubated on ice for 10 minutes. After incubation, cells were agitated using a P1000 pipette and 10 μL of sample was imaged. DAPI (Sigma, Cat #D9542) and Wheat Germ Agglutinin (WGA; Thermo Fisher Scientific, W11262) were used to determine if the cellular membrane had properly lysed for each cell. If intact cells were still present, 2 mL of sample was moved to a glass dounce tissue grinder (Sigma, Cat #D8938) and dounced 5 times. After douncing, another 10 μL sample was imaged under the microscope with DAPI and WGA staining as before to determine if high-quality, intact nuclei were obtained (see Figure 5B). We adjusted the number of dounces until only nuclei were found. As iPSCs differentiated further into CMs, the number of required dounces needed to be increased. For example, day 3 of differentiation required 5 dounces to obtain proper cell lysis and intact nuclei, while Day 7 required 12 dounces. Nuclei were spun down at 500 xg for 5 minutes at 4 °C. After centrifugation, the nuclei were washed with nuclei suspension buffer (NSB; 1X PBS, 0.01% BSA, and 0.1% RNAse inhibitor (Lucigen, #F83923)), resuspended in 2 mL NSB and filtered using a 35 μm cell strainer (Corning, #352235). 10 μL of nuclei suspension was sampled using a NI hemocytometer and the concentration adjusted to a final loading concentration of 300,000 nuclei/mL in NSB of which 2 mL was used for DroNc-seq.

### Microfluidic Co-encapsulation of Cells/Nuclei and Barcoded Beads

For **Drop-seq**, 2 mL of cells at 200,000 cells/mL in PBS-BSA was loaded in a 3 mL syringe (BD, #309657). A custom-built 90 μm Drop-seq microfluidic device (CAD file supplied separately) was used for droplet generation, creating droplets smaller than the standard Drop-seq protocol.^5^ We chose to use the 90 µm droplets because the effective concentration of cellular RNA in the 90 um drops is doubled, leading to better RNA capture, compared to 125 µm droplets used in Drop-seq. Indeed, we see an increase in RNA capture for cells of smaller size, such as iPSC. We note that the increase in capture efficiency often fails to translate to larger sized cells (∼15 μm), likely due to the higher concentrations of the lysed cell’s endogenous RNase and lysosomes, etc. in the drop. Cells at 200,000 cells/mL and ∼2,600,000 droplets/mL (droplet volume is ∼380 pL) amounts to a Poisson loading distribution with λ ≈ 0.076. DNA barcoded beads (ChemGenes, Macosko-2011-10(V+)) were washed, filtered, and suspended in Drop-seq lysis buffer, also at 200,000 beads/mL and kept in suspension under constant stirring using a magnetic tumble stirrer and flea magnet (V&P Scientific, VP 710 Series, VP 782N-3-150). Beads and cells were co-flowed into the device, each at 3 mL/hr, along with a surfactant-oil mix (BioRad, #1864006) at 12 mL/hr that was loaded into a 10 mL syringe (BD, #302995) and used as the outer carrier oil phase. Reverse emulsions droplets were generated at ∼3000 drops/sec and collected in two batches of 20 minutes each in 50 mL tubes (Genesee Scientific, #28-106). After collection, the standard Drop-seq protocol for bead recovery, washing, and reverse transcription was followed.^5^ After washes and DNaseI treatment as per Drop-seq protocol^5^, cDNA amplification was performed on 75,000 RNA-DNA barcode bead conjugates in a 96-well plate (Genesee Scientific, #24-302) loaded at 5000 beads per well, for a total of 15 wells and amplified for 15 PCR cycles using template switching.^5^ Post-PCR cleanup was performed by removing the STAMPs (Single Transcriptome Attached to Micro-Particles^5^) and pooling the supernatant from the wells together into a single 1.7 mL tube (Genesee Scientific, #22-281LR) along with 0.6X Ampure XP beads (Beckman Coulter, #A63880). After adding the Ampure beads to the PCR product, the tube was incubated at room temperature for 2 minutes on a thermomixer (Eppendorf Thermomixer C, #5382000023) set to 1250 rpm, and for another 2 minutes on bench for stationary incubation. Next, the tube was placed on a magnet, and 4X 80% ethanol washes were performed with 1 mL ethanol added in each wash. cDNA was eluted in 150 μL of water and the concentration and library size were measured using Qubit 3 fluorometer (Thermo Fisher) and BioAnalyzer High Sensitivity Chip (Agilent, #5067-4626). A BioAnalyzer trace is provided in Figure 5C as an example of the amplified transcriptome obtained from a Drop-seq run. 450 pg of the cDNA library was used in Nextera Library prep, instead of 650 pg as suggested in the Drop-seq protocol^5^ to obtain Nextera libraries between 300 – 600 bp.

For **DroNc-seq**, a 75 μm microfluidic device^9^ was used. 2 mL of nuclei at 300,000 nuclei/mL were loaded into a 3 mL syringe and flowed at 1.5 mL/hr. Barcoded beads were filtered with a 40 μm filter to select for smaller beads to prevent clogging events in the relatively smaller microfluidic channels. 2 mL of beads were suspended at 350,000 beads/mL in Drop-seq lysis buffer, loaded in a 3 mL syringe, kept suspended through a magnetic tumble stirrer, and flowed at 1.5 mL/hr, along with carrier oil-surfactant mix loaded in 10 mL syringe and flowed at 12 mL/hr. Droplets were generated at ∼4,500 drops/sec and collected in 50 mL tubes in two batches for 22 minutes each. After collection, the standard DroNc-seq protocol for bead recovery and reverse transcription was followed.^9^ cDNA amplification was performed on the STAMPs as above, for 15-20 wells at 5000 beads per well, for 15 PCR cycles. Cleanup was performed after removing the STAMPs and adding 0.6X Ampure XP beads (Beckman Coulter, #A63880) to the pooled supernatant followed by room temperature incubation for 2-minutes on an Eppendorf thermomixer set to 1250 rpm and another 2-minute stationary incubation. Tubes were placed on a magnet and beads were allowed to migrate prior to 4X washes in 80% ethanol. cDNA was eluted in 10 μL of water per well and DNA concentration was measured using a Qubit 3 fluorometer (Thermo Fisher). 650 pg of DNA was used in each Nextera reaction for fragmenting, tagging, and amplifying to create Nextera library. Nextera library size and concentrations were determined using a BioAnalyzer DNA High Sensitivity Chip (Agilent, #5067-4626).

### Nuclei Isolation from Adult Human Heart Tissue

Post-mortem human heart tissue was provided by the National Disease Research Interchange (NDRI). The sample (m, 68 yrs) had been stored at −80°C for 11 years before it was processed for DroNc-seq. The frozen tissue sample was weighed and cut with a scalpel and 32.8 mg of sample was processed, by mincing with the scalpel. The sample was placed into a glass dounce tissue grinder (Sigma, Cat #D8938) with 2 mL of ice-cold EZ-Prep lysis buffer from the Nuclei EZ-prep Isolation Kit (Sigma, Cat #NUC-101). The tissue was dounced 25 times with Pestle A, transferred to a conical tube with an additional 2 mL lysis buffer, and incubated on ice for 5 minutes. Sample was then centrifuged at 500 xg for 5 minutes at 4 °C. Supernatant was aspirated off and replaced with 2 mL lysis buffer. Sample was transferred back to the tissue grinder and dounced 25 times with Pestle B. Sample was then put back into a conical tube with an additional 2 mL lysis buffer, centrifuged, and washed with 4 mL lysis buffer followed by 5-minute incubation on ice. 10 μL of sample was taken and combined with DAPI and Wheat Germ Agglutinin (WGA) and put into an NI hemocytometer (InCyto, DHC-N01-2) to check for nuclei quality. If whole cells were still present, additional douncing with Pestle B was performed (additional 25-35 dounces expected) before checking again using DAPI and WGA. The resulting nuclei were centrifuged, lysis buffer was aspirated, and nuclei were washed and resuspended in Nuclei Suspension Buffer (NSB; 1x PBS, 0.01% BSA, and 0.1% RNAse inhibitor (Lucigen, #F83923)). Nuclei were filtered once with a 35 μm cell strainer (Corning, #352235), once with a 20 μm filter (pluriSelect, #43-50020-01), and twice with a 10 μm filter (pluriSelect, #43-50010-01) and stored on ice for processing. Nuclei were counted using an NI hemocytometer and brought to a final concentration of 300,000 nuclei/mL in 2 mL NSB for DroNc-seq. To assess the quality of RNA from the archived heart tissue, we ran an independent experiment to extract total RNA using a Qiagen kit (Qiagen, #74004) and measured using a BioAnalyzer RNA 6000 Pico kit (Agilent, #5067-1513). A RIN score of ∼5 was obtained for this sample.

### DroNc-seq on Nuclei Harvested from Heart Tissue

DroNc-seq was performed as previously described with a few exceptions: single 30-minute droplet collection was performed using a 75 μm microfluidic device and flow rates mentioned previously. During whole transcriptome amplification, 12 cycles of PCR were performed on 30 wells with 5000 barcoded beads per well. Clean-up was performed as described above. cDNA from each well was eluted in 2 μL of water and pooled for quantification by BioAnalyzer (Figure 5C) and Qubit, followed by Nextera library preparation.

### Sequencing

Drop-seq and DroNc-seq samples for each differentiation time-point were sequenced in a single run, with 150-200 million reads allocated per sample. Sample libraries were loaded at ∼1.5 pM concentration and sequenced on an Illumina NextSeq 500 using the NextSeq 75 cycle v3 kits for paired-end sequencing. 20 bp were sequenced for Read 1, 60 bp for Read 2 using Custom Read 1 primer, GCCTGTCCGCGGAAGCAGTGGTATCAACGCAGAGTAC^5^, according to manufacturer’s instructions. Illumina PhiX Control v3 Library was added at 5% of the total loading concentration for all sequencing runs.

### RNA-Seq Data Processing and Analyses

The differentiating iPSCs were sampled at specific timepoints during a 15-day period (days 0, 1, 3, 7, 15) using both Drop-seq and DroNc-seq (Fig 1A). A total of 17 sequencing runs were performed over the course of the differentiation. Each sequencing run produced paired-end reads, with one pair representing the 12 bp cell barcode and 8 bp unique molecular identifier (UMI), and the second pair representing a 60 bp mRNA fragment. We developed a Snakemake^21^ protocol that takes a FASTQ file with such paired-end reads as input and produces an expression matrix corresponding to the UMI of each gene in each cell. The protocol initially performs *FastQC*^22^ to obtain a report of read quality. Next, it creates a whitelist of cell barcodes using *umi_tools*^23^ *0.5.3*, which is a list of cell barcodes with at least 30k reads. Next, each paired-end read is combined into a single read where the read name contains the cell barcode and UMI extracted from paired end read 1, and the sequence content corresponds to paired end read 2. This is done for every paired end read and placed into a single “tagged” FASTQ file. The tagged FASTQ file contains only the cell barcodes found in the whitelist. Finally, the protocol trims the ends of reads to remove polyA sequences and adaptors using *cutadapt*^24^ *1.15*. The tagged and trimmed FASTQ file is aligned to the human reference genome (version GRCh38) using the *STAR*^25^ aligner version 2.5.3, which returns a BAM file sorted by coordinate. Next, we use *featureCounts*^26^ version 1.6.0 to assign each aligned read to a feature on the genome. Finally, we use the *count* function from *umi_tools* to create a count matrix representing the frequency of each feature in the BAM file. The pipeline is available at github.com/aselewa/dropseq_pipeline. A total of 17 count matrices were produced by this pipeline, 9 of which correspond to Drop-seq and 8 correspond to DroNc-seq. In order to incorporate introns into the counting process, the UMI count of a gene was calculated as the sum of its exon and intron UMIs. This is particularly important for DroNc-seq as approximately half the reads obtained come from intronic regions of pre-spliced mRNA. GENCODE version 28 annotations contain exon features and gene features but do not contain intron features. To derive an intron annotation file, we used exon and gene features. Exon regions were subtracted from gene regions (on the same strand) and the remainder was counted as the intron region for said gene. Then the expression level of a gene is given by the sum of the number of intron and exons.

From each sequencing run, approximately 5000 cells were obtained with an average read depth of 30k – 40k per cell. Low quality cells were filtered based on the number of genes detected. A gene was considered detected in a cell if there was at least 1 UMI present. Cells with less than 400 genes and nuclei with less than 300 genes detected were removed. Low quality genes were also filtered if they were not detected in at least 10 cells, in order to reduce noise and computation cost. The total numbers of cells remaining were approximately 23,554 and 24,318 for Drop-seq and DroNc-seq, respectively. After filtering, all expression matrices from Drop-seq experiments were merged into a single expression matrix. The merging was done by taking the union of all genes. If a particular dataset did not contain a gene that is expressed in another dataset, we set the expression level to zero in the first dataset. Similarly, all expression matrices corresponding to DroNc-seq were merged into a single expression matrix. Both merged matrices were processed and analyzed separately downstream. Seurat^15^ was used to perform normalization, clustering, and cell type analysis. R scripts used for the analyses in this paper are documented at github.com/aselewa/czi.

### Internal Priming

We used the MEME^27^ suite to find all 5 bp stretches of adenines using the human genome build hg38. Next, we merged all 5 bp motifs in order to obtain all continuous polyA tracts. A total of ∼2 × 10^7^ motifs at least 5 bp long were identified genome-wide. BAM files from each time-point were merged and only intronic reads were kept. Intronic reads were extended by 20 bp on each side and intersected with the adenine motifs in a strand-specific way. The motifs were centered by the coordinates of the reads they intersect with and a histogram motif of 3’ positions was obtained (Figure S3).

### Normalization and Scaling

Following the analysis procedure recommended by Seurat, we first normalize the count data. Each cell’s gene-specific UMIs were divided by the total number of UMI in the cell scaled to 10^4^, which yields TP10k (transcripts per 10k) values. Figure S2 shows the relationship between the mean expression (mean TP10k) and the length of the gene. The relationship is relatively weak, therefore normalizing by just the library size is sufficient. A pseudo-count of 1 was added to all scaled values followed by a natural log transformation. After the log-transformation, the values were standardized, i.e. mean-centered and scaled such that each gene has unit variance. These log-normalized, and standardized data were used in downstream analyses to perform dimensionality reduction and reconstruction of differentiation trajectories.

### Dimensionality Reduction

The first step performed in dimensionality reduction is principal components analysis (PCA). Prior to PCA, Seurat calculates the gene dispersion vs. mean expression in order to obtain a subset of highly variable genes, which reduces the computational time of PCA compared with using the entire subset of genes identified in the experiment. Highly variable genes were selected based on a threshold of 1.5 for the dispersion level and a minimum expression level of 0.15 (on log scale) yielding 400 genes and 350 genes with Drop-seq and DroNc-seq, respectively. These highly variable genes were used to calculate principal components for Drop-seq and DroNc-seq data. The top 7 principal components, which explained 60% and 70% of variation for Drop-seq and DroNc-seq, respectively, were used to perform clustering and the results were visualized with the Uniform Manifold Approximation and Projection (UMAP^28^), which produced a 2-dimensional visualization of the data (Figures S4 A, B left). We also performed tSNE on the same data (Figures S4 A, B right) using a perplexity of 50 and found that UMAP captures more of the global structure in the data, as previously reported^29^. A minimum distance of 0.5 and 0.6 were used in UMAP for Drop-seq and DroNc-seq, respectively.

### Cell type Analysis

The principal components were used for graphical clustering using the *FindClusters* command of Seurat. A resolution parameter of 0.13 is used to obtain 6 clusters in Drop-seq and 5 clusters in DroNc-seq. In order to determine cell types from the clusters, we performed differential expression analysis using the *FindAllMarkers* function and *negbinom* test in Seurat. This identifies differentially expressed genes between every two groups of cells using a likelihood ratio test of negative binomial generalized linear models. The Seurat’s *negbinom* test yields relatively low false positive rates for differential expression analyses, compared with other parametric methods^30^. The p-values were adjusted for multiple testing using the Bonferroni correction. Furthermore, as we were only interested in upregulated genes as these will define the cell type, we ordered genes in each cluster, by their average log-fold-change (logFC) in descending order. Marker genes were identified based on functional annotations as these genes associated with cell types have a large fold-change in expression. Figures S6 and S7 show the top 10 differentially expressed genes in each identified cluster for Drop-seq and DroNc-seq, respectively.

### Pseudo-bulk Analysis

Raw RNA-seq counts were obtained from GEO accession GSE110471 and the human samples were extracted from the population. The raw counts were converted into log-TP10k’s. After filtering low-quality cells, Drop-seq and DroNc-seq counts were aggregated (summed) for each gene, and the resulting counts were converted to log (TP10k + 1). The Pearson correlation between pseudo-bulk and each bulk RNA-seq sample was calculated using the *cor* function in *R 3.5.1* across ∼6,000 genes.

### Single-cell Trajectory Analysis

*Monocle* version 2.6.4 was used to construct single-cell differentiation trajectories. Computing the trajectory of approximately 20,000 cells is computationally expensive and slow with Monocle. To overcome this, we used the best 700 cells from each time-point. In particular, cells were ordered by their detection rate (number of genes detected) and 700 cells with the highest detection rate were chosen. The computation is also expensive and slow when the number of genes is high (>10,000 genes). Selection of genes for trajectory analysis, or feature selection, is critical for obtaining accurate trajectories. In our case, we used all of the differentially expressed genes in the cell type analysis. The data given to Monocle are log-transformed TP10k values. The *reduceDimension* function with the *DDRTree* method was used to obtain a 2-dimensional representation of the developmental trajectories in each dataset. The cells were then ordered using the *orderCell*s function, which infers the trajectory in reduced-dimension dataset using reserve graph embedding^17^. A total of 3,500 cells (700 per time-point, 5 time-points in total) were used to infer the trajectories in Drop-seq and DroNc-seq.

### Primary Heart Tissue Analysis

A total of 4796 nuclei obtained from post-mortem adult human male heart tissue were profiled using DroNc-seq. Genes were quantified using both introns and exons, with mean number of genes and UMIs of 361 and 823, respectively. The top 30% of cells were chosen based on number of genes detected, which corresponds to 1,491 cells. Transformation of data and cell type analysis was performed in the manner described above. Next, we calculated the Pearson correlation coefficient using the *cor* function in *R 3.5.1* between primary heart nuclei and the *in vitro* iPSC-derived CMs profiled by DroNc-seq. We also used iPSCs profiled by DroNc-seq as an out-group. A total of 200 iPSC-derived CMs and 50 iPSCs were used for the correlation analysis. For each primary heart nuclei, a total of 250 correlation coefficients were calculated using ∼2500 genes, which we call the correlation profile of a cell. The resulting matrix of correlation values were visualized and bi-clustered with the *heatmap*. 2 function in *R 3.5.1.*

## Data Availability

All raw data are available through the Human Cell Atlas Portal (https://prod.data.humancellatlas.org/explore/projects/c765e3f9-7cfc-4501-8832-79e5f7abd321). All code used for analysis is available at github.com/aselewa/czi and github.com/aselewa/dropseq_pipeline.

## Acknowledgement

We thank Megan Rowton, Alex Guzzetta, and John Blischak for helpful comments on the manuscript. This work was supported by the Chan-Zuckerberg Initiative pilot award #2017-174052. RE was supported by the NIH MSTP Training Grant T32GM007281. KR was supported by NIH GRTG 5T32GM007197 and AHA Predoctoral Fellowship 18PRE34030197. SP was supported by the National Center for Advancing Translational Sciences of the NIH (K12 HL119995). This work was performed, in part, at the Center for Nanoscale Materials, a U.S. Department of Energy Office of Science User Facility, and supported by the U.S. Department of Energy, Office of Science, under Contract No. DE-AC02-06CH11357.

## Supplementary Methods

### Species-mixing and single-cell specificity

For the Drop-seq experiment on biological replicate #1, chimpanzee iPSCs^20^ were mixed with human iPSC-derived CMs from day 7 of the differentiation time-point, in order to assess the frequency of doublets during cell encapsulation. We used chimpanzee cells for the species mixing as these cells were grown using identical conditions as the human cells. The alignment protocol was adjusted so that each read was aligned to both the human genome (GRCh38) and the chimp genome (panTro5) separately. For each cell that passed quality control, we counted the number of reads that aligned exclusively or with a better score to the genome of one of the species (Figure S1). We then used the ratio of these counts as a ‘species-specificity’ score for each cell. We found only a small number of cells with scores that could suggest mixing of cells from human and chimp (< 5%), similar to previously reported estimates^5^. Cells with intermediate scores had typically lower read counts and were thus removed by filtering based on read depth. We only kept cells with a specificity score above 0.6 yielding ∼739 cells. In agreement with our assignment, > 99% of these cells were associated with clusters that we identified as CMs while none were associated with iPSC clusters.

**Figure S1:**
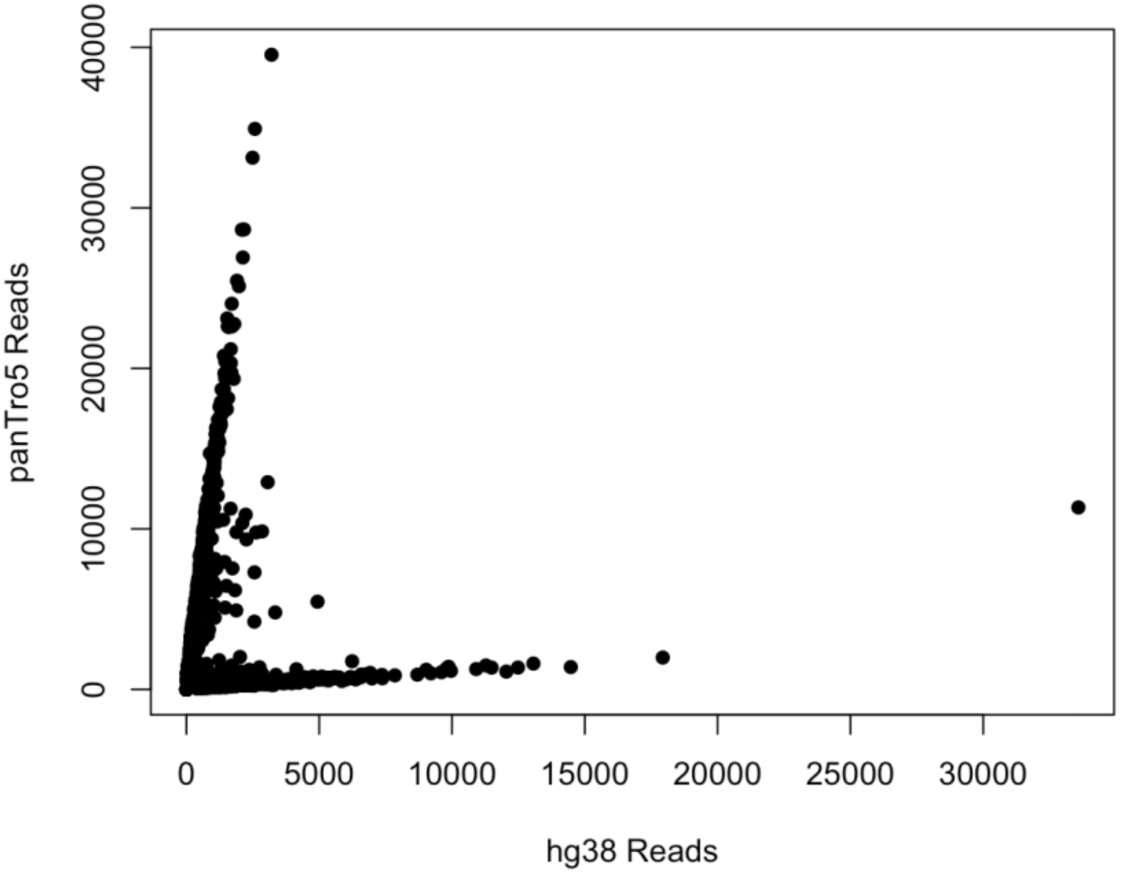
Scatterplot of number of reads assigned to hg38 vs panTro5 for each cell in Drop-seq day 7, cell line #1 as part of a species-mixing experiment using human iPSC derived cardiomyocytes and chimpanzee iPSCs.

**Figure S2:**
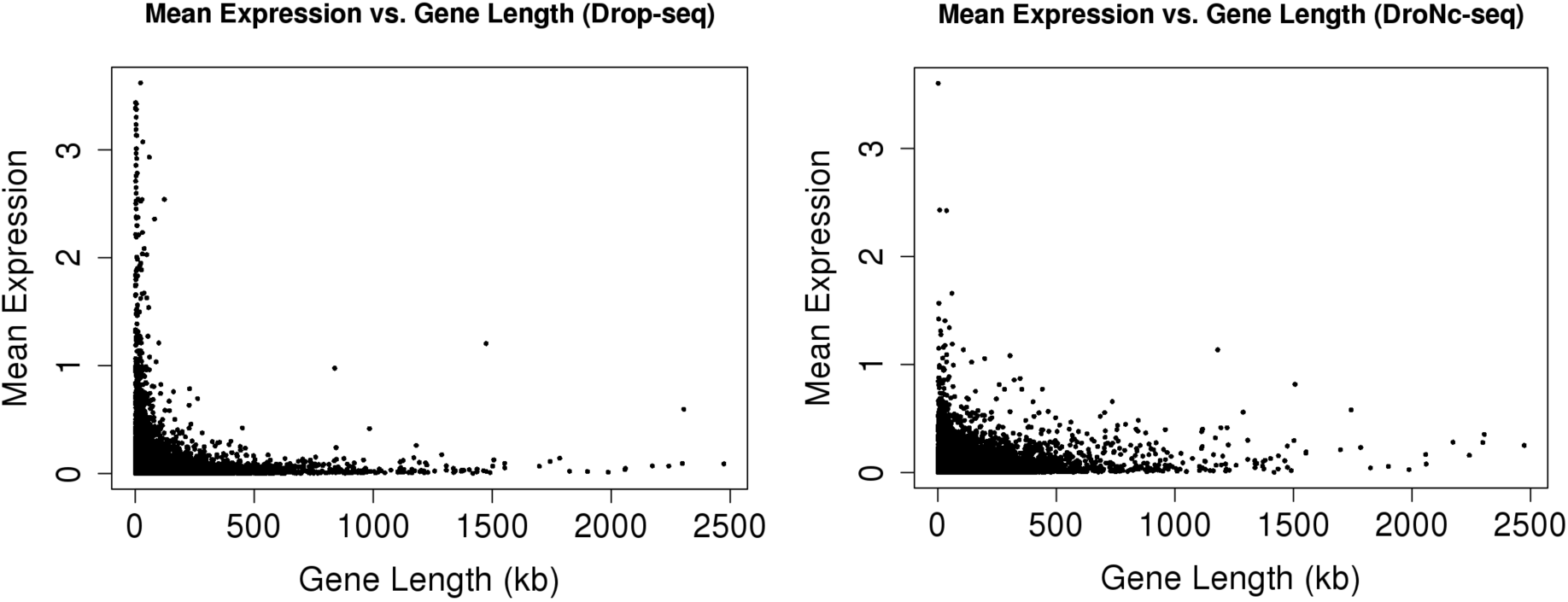
Mean expression (log) vs. gene length for Drop-seq (left) and DroNc-seq (right).

**Figure S3:**
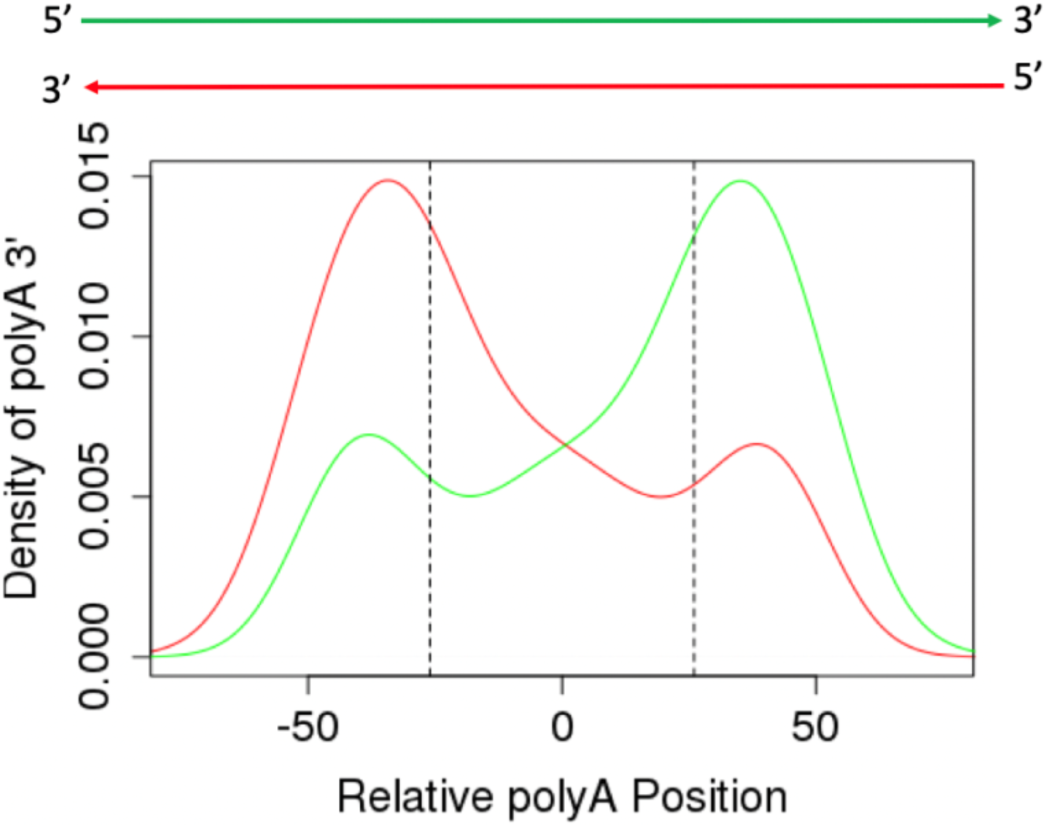
Density curves of the position of polyA at the 3’ end. Green and red curves represent reads mapping to the forward and reserve direction, respectively. The dashed line represents the average read length.

**Figure S4:**
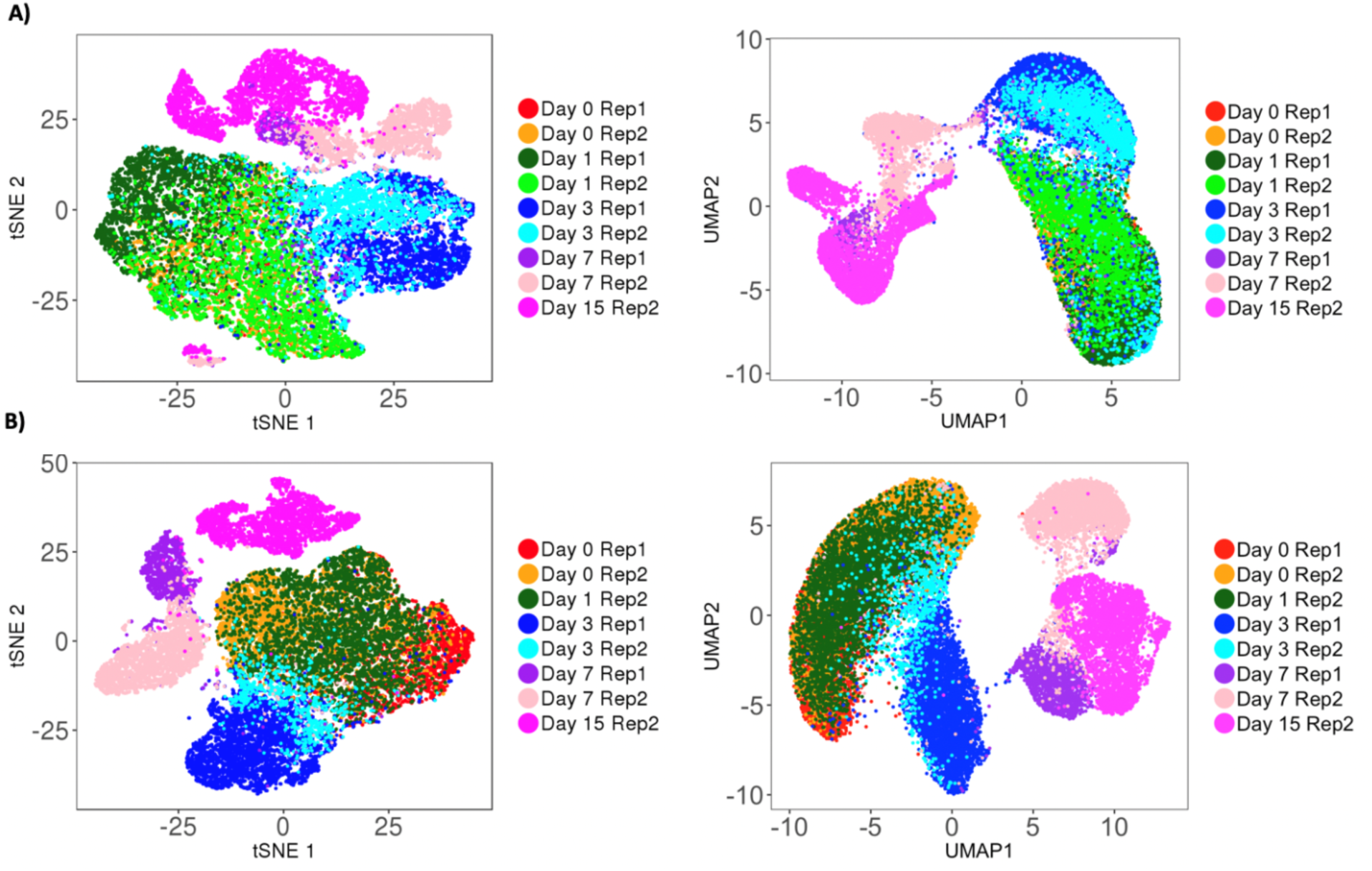
Dimensionality reduction for A) Drop-seq and B) DroNc-seq using tSNE (left) and UMAP (right). Color represents the differentiation time point.

**Figure S5:**
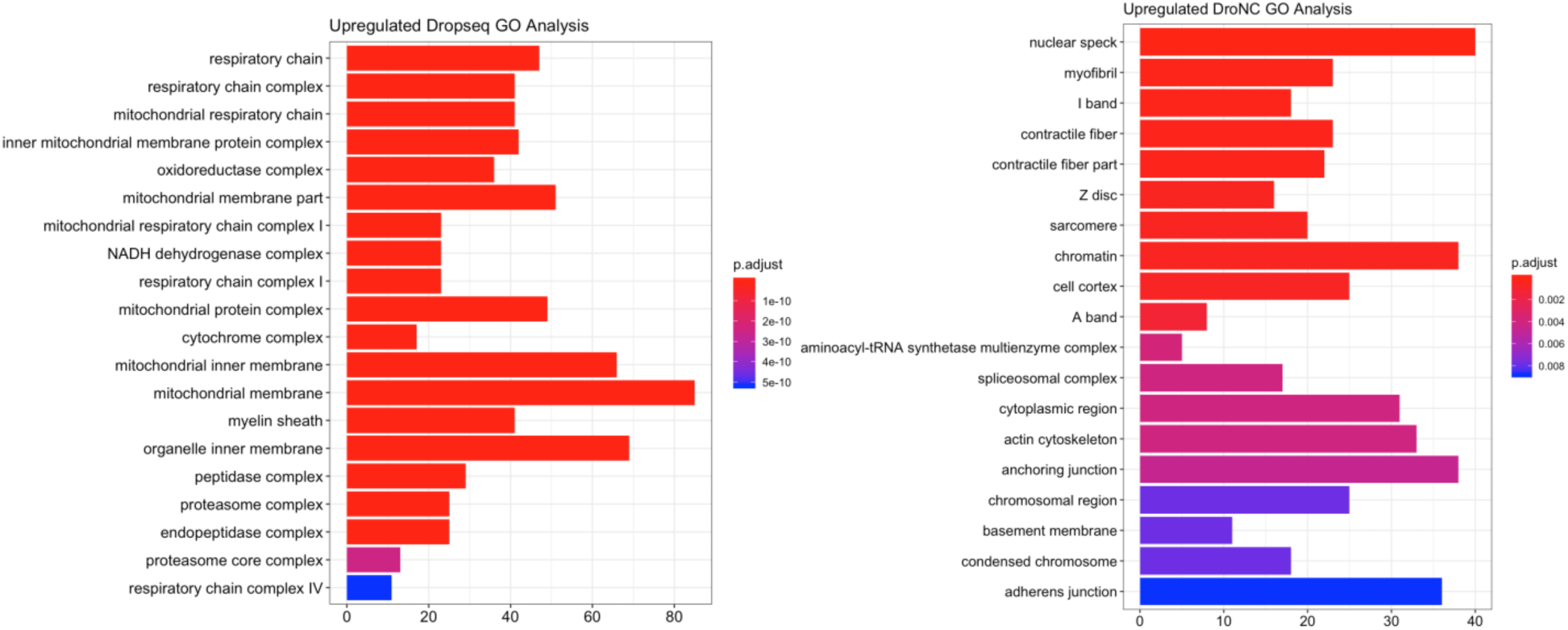
Gene enrichment analysis on differentially expressed genes between Drop-seq and DroNc-seq.

**Figure S6:**
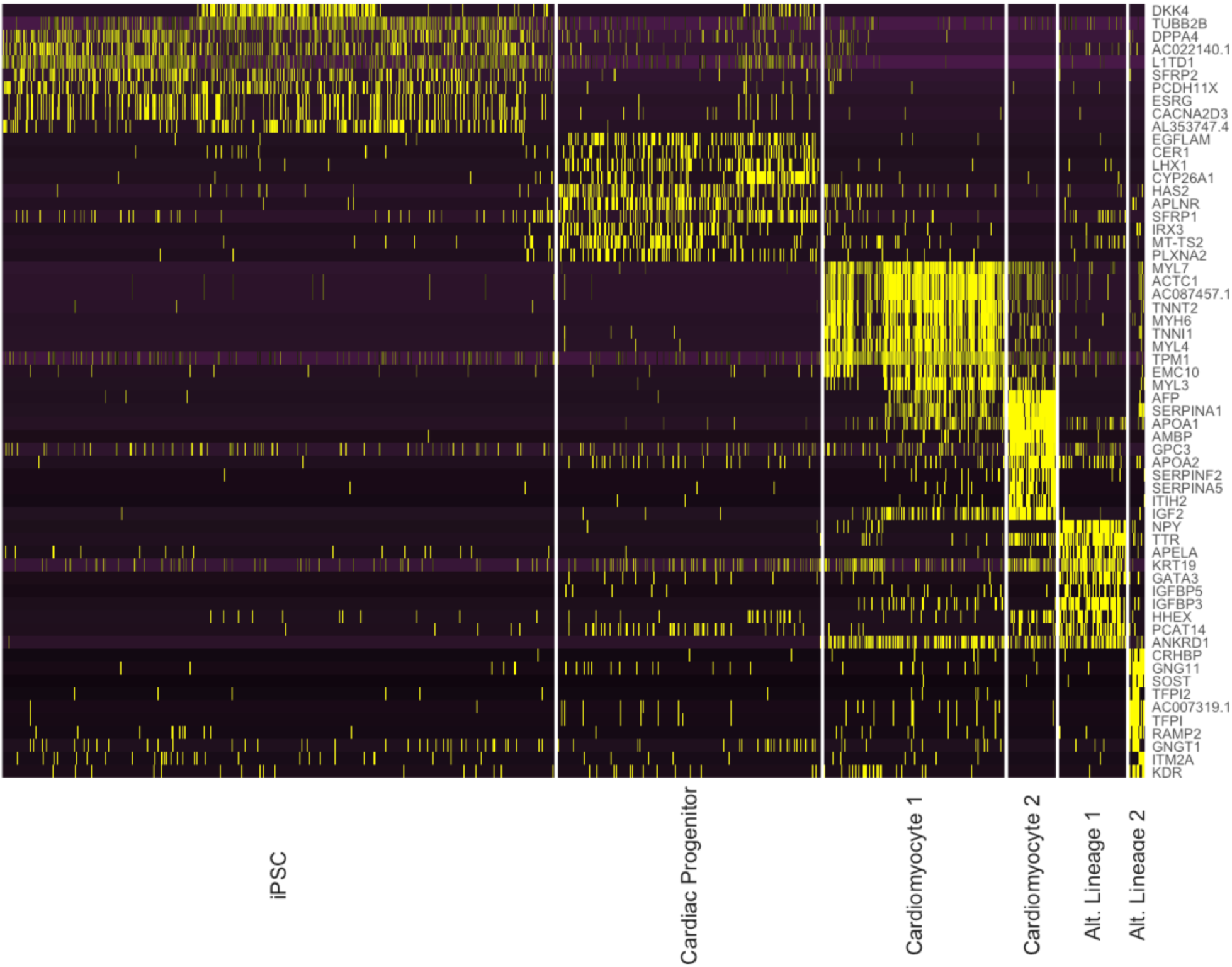
Heatmap of expression values of top 10 differentially expressed genes in each cell type cluster for Drop-seq.

**Figure S7:**
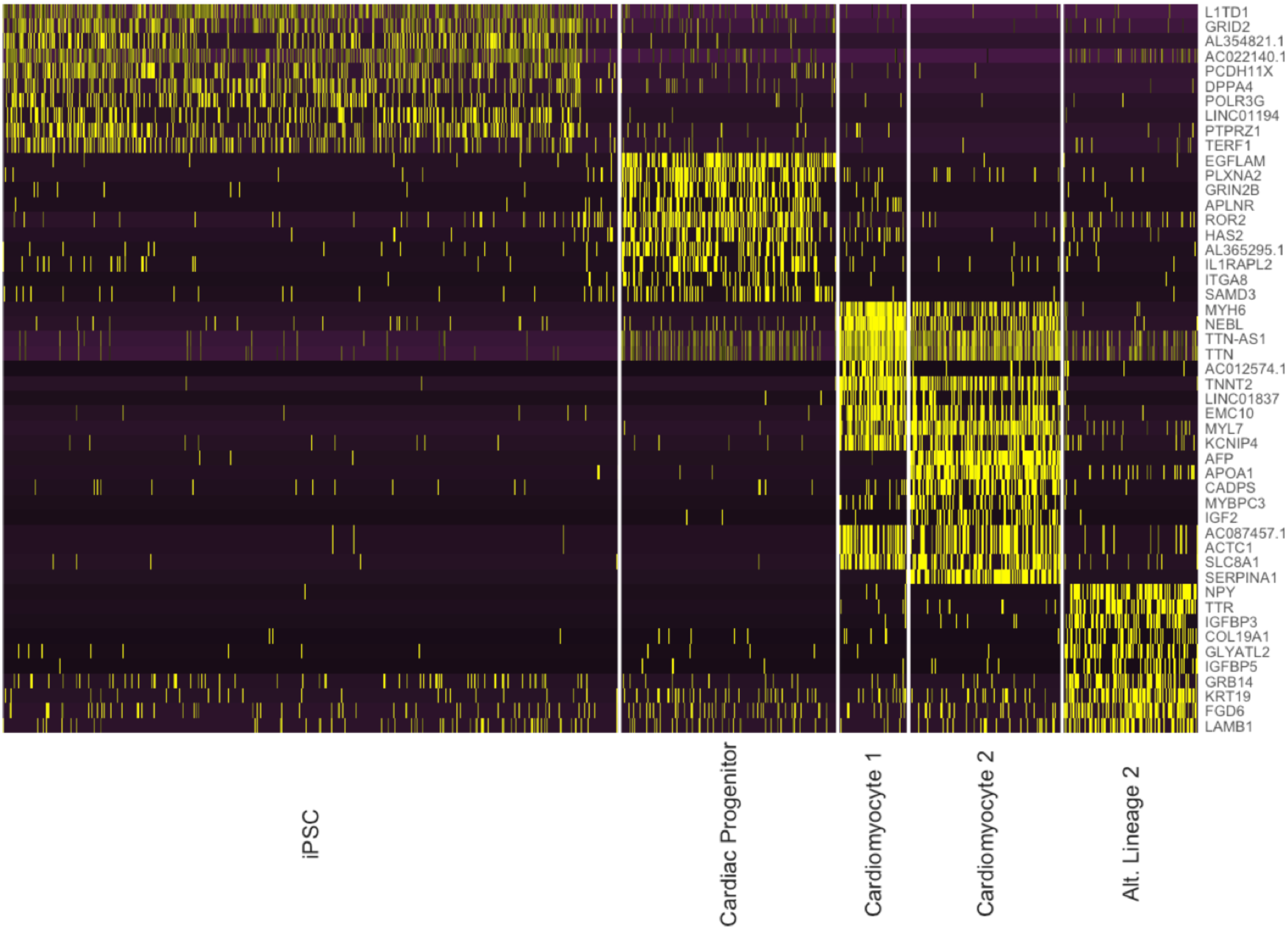
Heatmap of expression values of top 10 differentially expressed genes in each cell type cluster for DroNc-seq.

**Table S1:** Breakdown of cell types and associated genes discovered in Drop-seq and DroNc-seq

| Markers | Cell type | Prevalence (Drop-seq) | Prevalence (DroNc-seq) | Drop-seq Only Genes (top 5) | DroNc-seq Only Genes (top 5) |
| --- | --- | --- | --- | --- | --- |
| DPPA4 | iPSC | 48.9% | 52% | SFRP2, AC025465.1, ESRG, CACNAD2D3, BDNF-AS | RIMS2, RPL8, GOLGA4, EIF4A2, SET |
| EOMES<br>APLNR | Cardiac Progenitor | 23.3% | 18.2% | CER1, LHX1, CYP26A1, IRX3, MT-TS2 | GRIB2B, AL3365295.1, IL1RAPL2, KCNQ5, NRX3 |
| MYH6<br>TNNT2 | Cardiomyocyte 1 | 16.1% | 5.6% | MYL3, NPPA-AS1, NPPA, | AC012574.1, AC105233.5, MYO1D, |
|  |  |  |  | ACTN2,<br>TNNC1 | ARHGAP42,<br>CDK14 |
| MYH6<br>TNNT2<br>AFP<br>SERPINA1 | Cardiomyocyte 2 | 4.2% | 12.7% | AMBP,<br>APOA2,<br>SERPINF2,<br>ITIH2,<br>SERPINA5 | KCNH7,<br>ERBB4,<br>ZBTB20,<br>NRG3, KCNQ5 |
| TTR<br>FOXA2 | Alternative<br>Lineage 1 | 5.9% | 11.3% | GATA3,<br>S100A14,<br>HHEX,<br>FLIRT3,<br>EPSTIL1 | EWSR1,<br>PTBP2,<br>ZMYM2,<br>LUC7L,<br>LINC01876 |
| CD34<br>SCARF1<br>FLT1 | Alternative<br>Lineage 2 | 1.4% | 0% | CRHBP,<br>GNG11,<br>SOST, TFPI2,<br>AC007319.1 | None |

**Figure S8:**
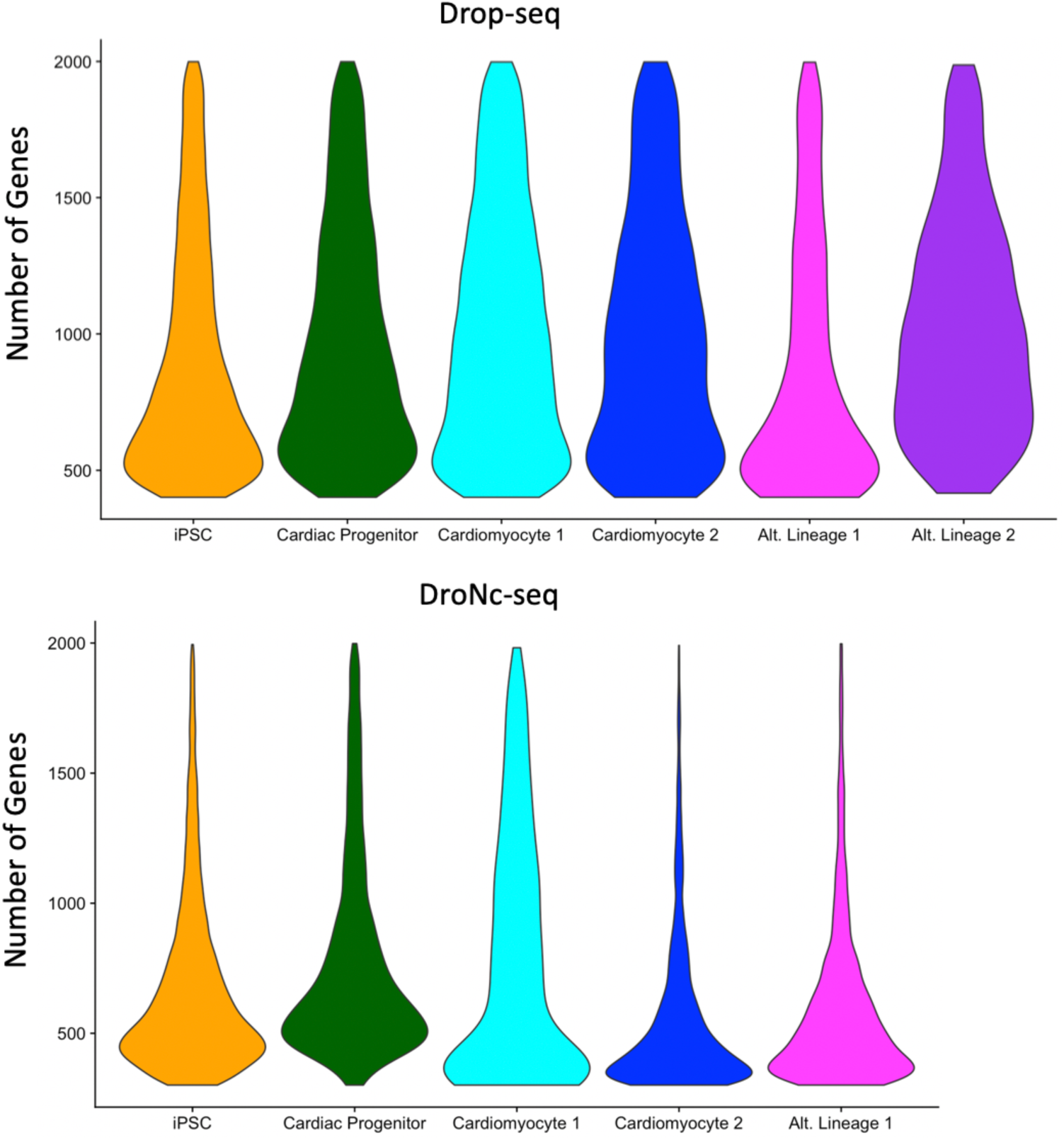
Violin plots representing the of number of genes in each cluster for Drop-seq (top) and DroNc-seq (bottom).

**Figure S9:**
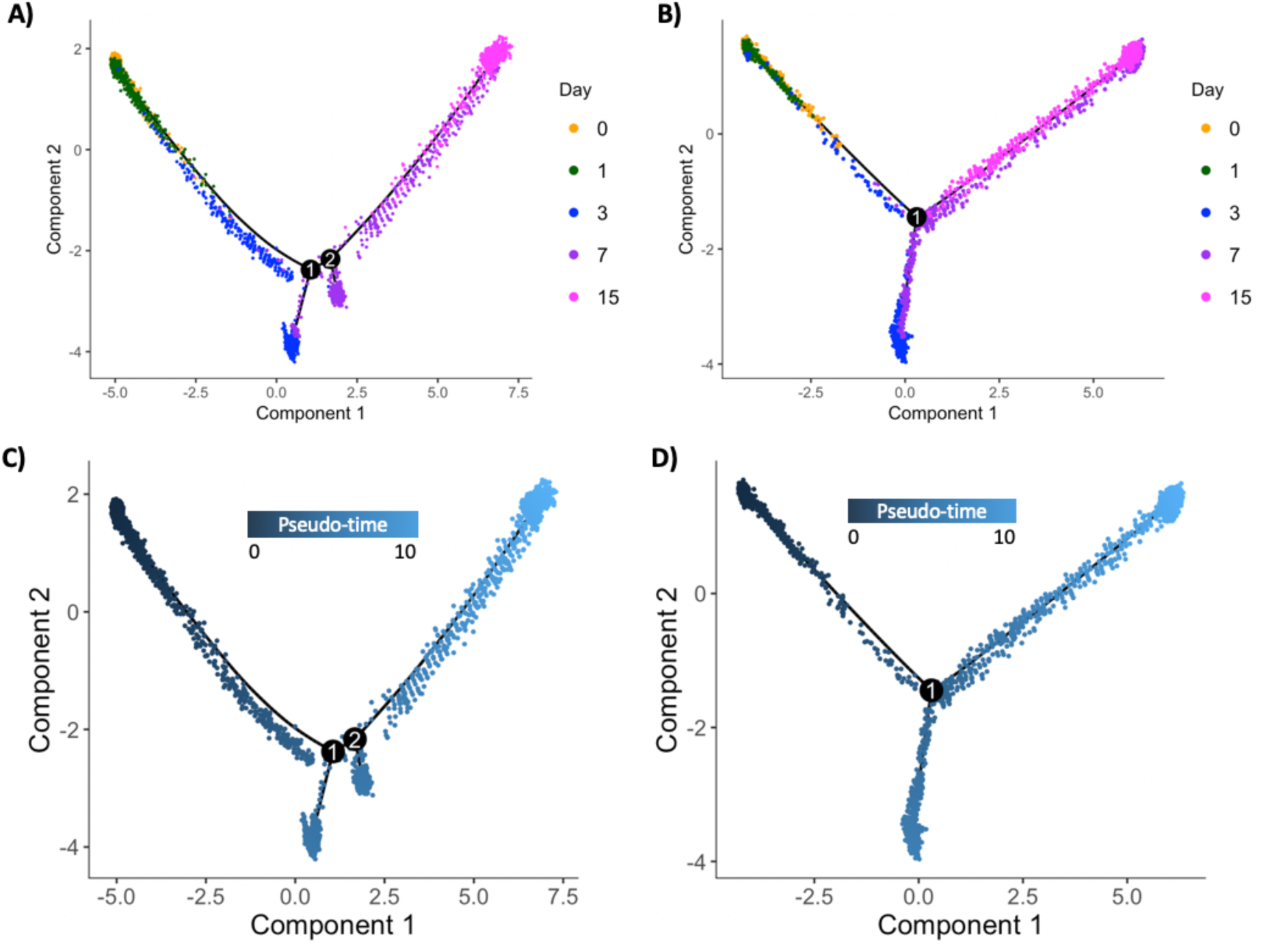
Cell differentiation trajectories constructed from Drop-seq (left), and DroNc-seq (right) using Monocle. Each differentiation time-point sampled is labelled by the same color in both techniques. A, B) uses the time-point as color, and C, D) shows the inferred pseudo-time as the color.

**Figure S10:**
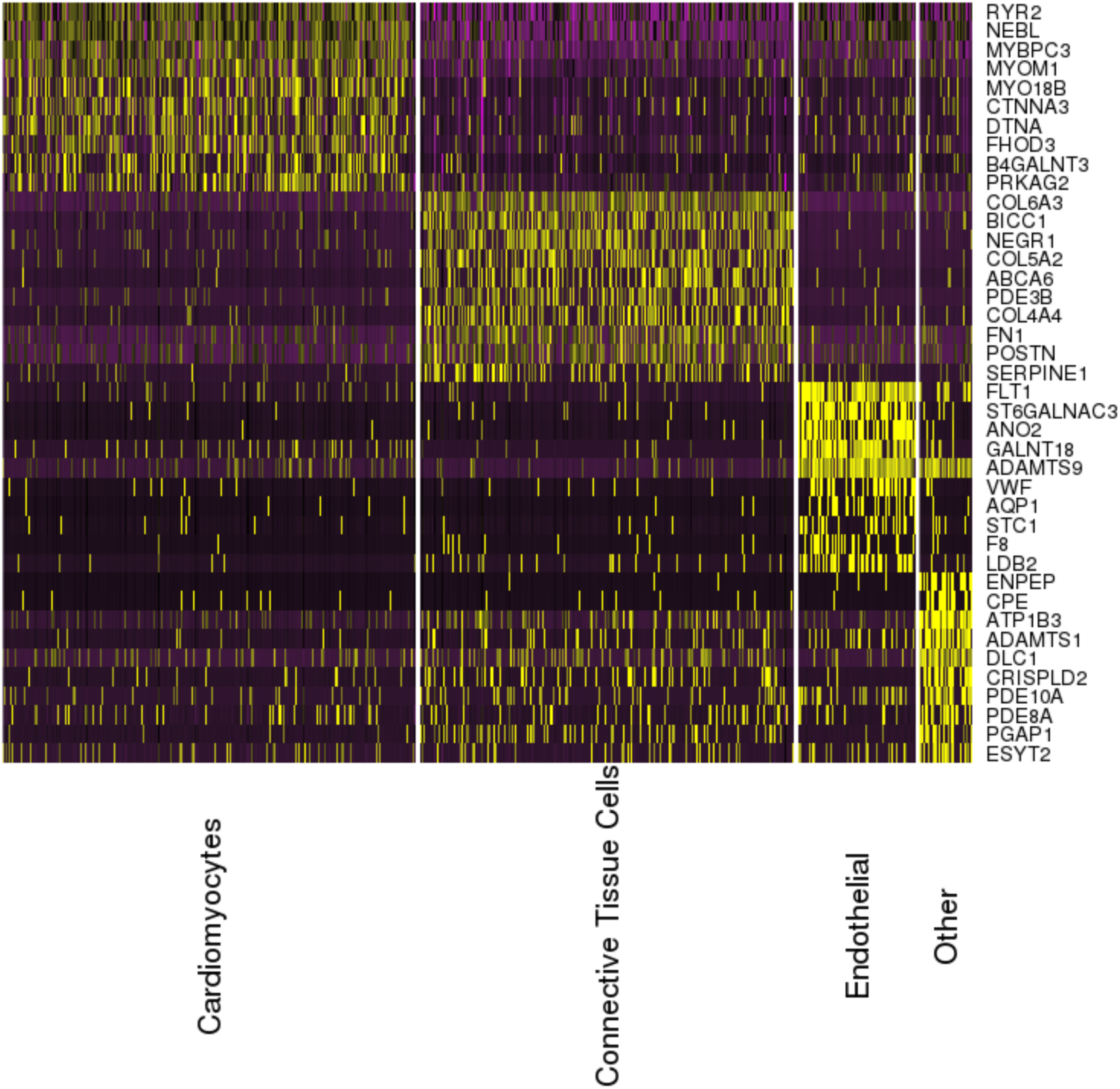
Top 10 upregulated genes identified in each cell type cluster using DroNc-seq on primary tissue from archived adult human heart.

**Figure S11:**
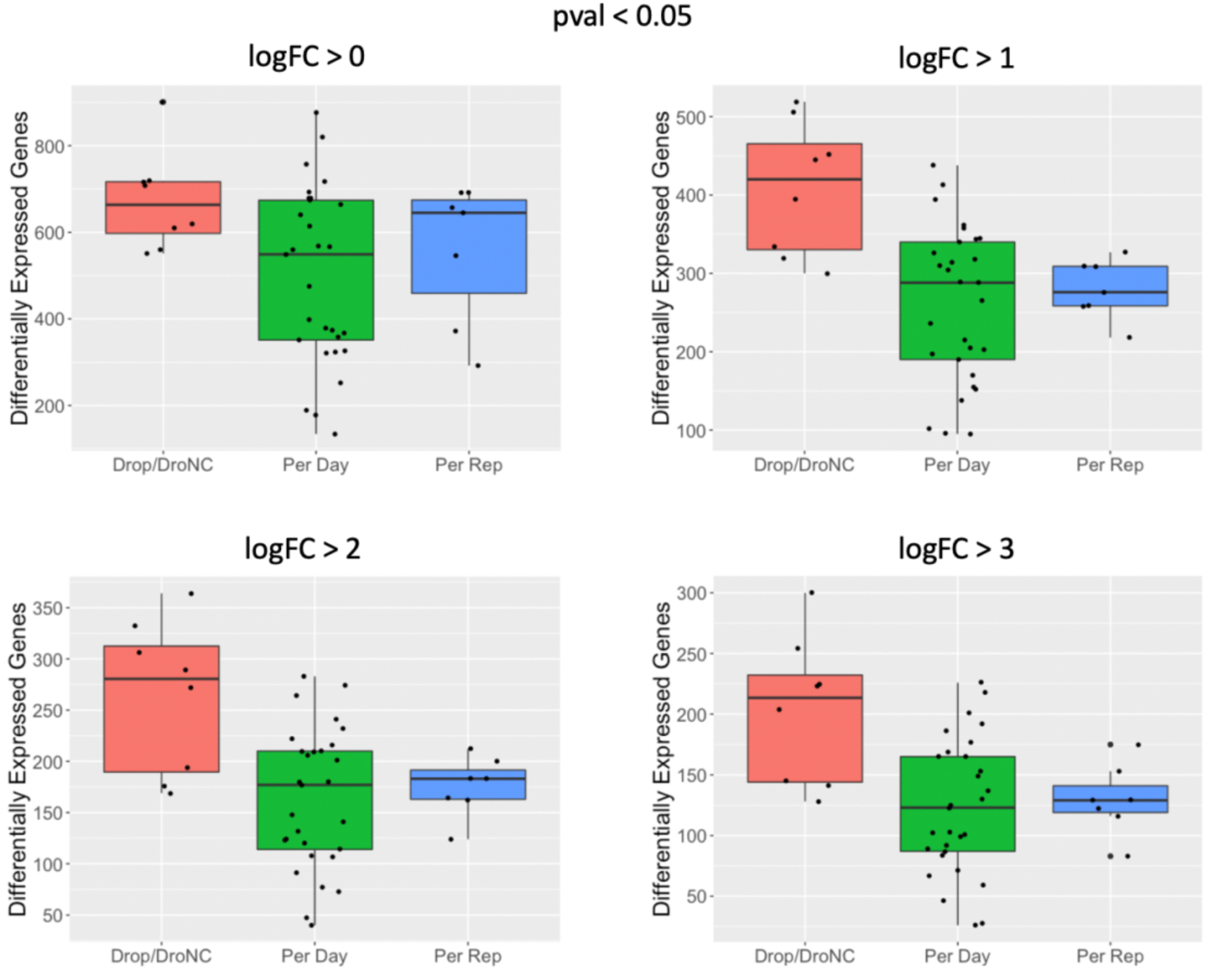
Differential expression analysis across time-points, cell-lines (biological replicates), and across Drop-seq and DroNc-seq using different thresholds for log-fold-change. All genes shown have adjusted p-value < 0.05.

## References

1. Rozenblatt-Rosen, O., Stubbington, M. J. T., Regev, A. & Teichmann, S. A. The Human Cell Atlas: From vision to reality. Nature (2017). doi:10.1038/550451a

2. Jaitin, D. A. et al. Massively parallel single-cell RNA-seq for marker-free decomposition of tissues into cell types. Science (80-.). (2014). doi:10.1126/science.1247651

3. Shalek, A. K. et al. Single-cell transcriptomics reveals bimodality in expression and splicing in immune cells. Nature (2013). doi:10.1038/nature12172

4. Treutlein, B. et al. Reconstructing lineage hierarchies of the distal lung epithelium using single-cell RNA-seq. Nature (2014). doi:10.1038/nature13173

5. Macosko, E. Z. et al. Highly parallel genome-wide expression profiling of individual cells using nanoliter droplets. Cell 161, 1202–1214 (2015).

6. Klein, A. M. et al. Droplet Barcoding for Single-Cell Transcriptomics Applied to Embryonic Stem Cells Accession Numbers GSE65525 Klein et al Resource Droplet Barcoding for Single-Cell Transcriptomics Applied to Embryonic Stem Cells. Cell (2015). doi:10.1016/j.cell.2015.04.044

7. Poran, A. et al. Single-cell RNA sequencing reveals a signature of sexual commitment in malaria parasites. Nature (2017). doi:10.1038/nature24280

8. Karaiskos, N. et al. The Drosophila embryo at single-cell transcriptome resolution. Science (80-.). (2017). doi:10.1126/science.aan3235

9. Habib, N. et al. Massively parallel single-nucleus RNA-seq with DroNc-seq. Nat. Methods 14, 955–958 (2017).

10. Lake, B. B. et al. A comparative strategy for single-nucleus and single-cell transcriptomes confirms accuracy in predicted cell-type expression from nuclear RNA. Sci. Rep. (2017). doi:10.1038/s41598-017-04426-w

11. Bakken, T. E. et al. Single-nucleus and single-cell transcriptomes compared in matched cortical cell types. PLoS One (2018). doi:10.1371/journal.pone.0209648

12. Wu, H., Kirita, Y., Donnelly, E. L. & Humphreys, B. D. Advantages of Single-Nucleus over Single-Cell RNA Sequencing of Adult Kidney: Rare Cell Types and Novel Cell States Revealed in Fibrosis. J. Am. Soc. Nephrol. (2018). doi:10.1681/asn.2018090912

13. Banovich, N. E. et al. Impact of regulatory variation across human iPSCs and differentiated cells. Genome Res. 28, 1243–1252 (2017).

14. La Manno, G. et al. RNA velocity of single cells. Nature (2018). doi:10.1038/s41586-018-0414-6

15. Butler, A., Hoffman, P., Smibert, P., Papalexi, E. & Satija, R. Integrating single-cell transcriptomic data across different conditions, technologies, and species. Nat. Biotechnol. (2018). doi:10.1038/nbt.4096

16. Friedman, C. E. et al. Single-Cell Transcriptomic Analysis of Cardiac Differentiation from Human PSCs Reveals HOPX-Dependent Cardiomyocyte Maturation. Cell Stem Cell (2018). doi:10.1016/j.stem.2018.09.009

17. Pavlovic, B. J., Blake, L. E., Roux, J., Chavarria, C. & Gilad, Y. A Comparative Assessment of Human and Chimpanzee iPSC-derived Cardiomyocytes with Primary Heart Tissues. Sci. Rep. (2018). doi:10.1038/s41598-018-33478-9

18. Setty, M. et al. Wishbone identifies bifurcating developmental trajectories from single-cell data. Nat. Biotechnol. (2016). doi:10.1038/nbt.3569

19. Trapnell, C. et al. The dynamics and regulators of cell fate decisions are revealed by pseudotemporal ordering of single cells. Nat. Biotechnol. (2014). doi:10.1038/nbt.2859

20. Romero, I. G. et al. A panel of induced pluripotent stem cells from chimpanzees: A resource for comparative functional genomics. Elife (2015). doi:10.7554/eLife.07103.001

21. Köster, J. & Rahmann, S. Snakemake-a scalable bioinformatics workflow engine. Bioinformatics (2012). doi:10.1093/bioinformatics/bts480

22. Andrews, S. & Babraham Bioinformatics. FastQC: A quality control tool for high throughput sequence data. Manual (2010). doi:citeulike-article-id:11583827

23. Smith, T., Heger, A. & Sudbery, I. UMI-tools: Modeling sequencing errors in Unique Molecular Identifiers to improve quantification accuracy. Genome Res. (2017). doi:10.1101/gr.209601.116

24. Martin, M. Cutadapt removes adapter sequences from high-throughput sequencing reads. EMBnet.journal (2011). doi:10.14806/ej.17.1.200

25. Dobin, A. et al. STAR: Ultrafast universal RNA-seq aligner. Bioinformatics (2013). doi:10.1093/bioinformatics/bts635

26. Liao, Y., Smyth, G. K. & Shi, W. FeatureCounts: An efficient general purpose program for assigning sequence reads to genomic features. Bioinformatics (2014). doi:10.1093/bioinformatics/btt656

27. Bailey, T. L. et al. MEME Suite: Tools for motif discovery and searching. Nucleic Acids Res. (2009). doi:10.1093/nar/gkp335

28. McInnes, L., Healy, J., Saul, N. & Großberger, L. UMAP: Uniform Manifold Approximation and Projection. J. Open Source Softw. (2018). doi:10.21105/joss.00861

29. Etienne, B. et al. Evaluation of UMAP as an alternative to t-SNE for single-cell data. Development (2018). doi:10.1101/298430

30. Soneson, C. & Robinson, M. D. Bias, robustness and scalability in single-cell differential expression analysis. Nat. Methods (2018). doi:10.1038/nmeth.4612

